# Abundant deep ocean heterotrophic bacteria are culturable

**DOI:** 10.1101/2022.12.16.520132

**Authors:** Isabel Sanz-Sáez, Pablo Sánchez, Guillem Salazar, Shinichi Sunagawa, Colomban de Vargas, Chris Bowler, Matthew B. Sullivan, Patrick Wincker, Eric Karsenti, Carlos Pedrós-Alió, Susana Agustí, Takashi Gojobori, Carlos M. Duarte, Josep M. Gasol, Olga Sánchez, Silvia G. Acinas

**Affiliations:** Departament de Biologia Marina i Oceanografia, Institut de Ciències del Mar, ICM-CSIC, Barcelona, Spain; Department of Biology, Institute of Microbiology, ETH Zurich, Vladimir-Prelog-Weg 1-5/10, CH-8093 Zurich, Switzerland; Sorbonne University, CNRS, Station Biologique de Roscoff, UMR7144, ECOMAP, Roscoff, France; Institut de biologie de l’Ecole normale supérieure (IBENS), Ecole normale supérieure, CNRS, INSERM, PSL Université Paris, 75005 Paris, France; Departments of Microbiology and Civil, Environmental and Geodetic Engineering; The Ohio State University; Columbus, OH 43210; USA; Génomique Métabolique, Genoscope, Institut de biologie François Jacob, Commissariat à l’Energie Atomique (CEA), CNRS, Université Evry, Université Paris-Saclay, Evry, France; Research Federation for the Study of Global Ocean Systems Ecology and Evolution, FR2022/Tara Oceans GOSEE, 75016 Paris, France; Institut de Biologie de l’Ecole Normale Supérieure, Ecole Normale Supérieure, CNRS, INSERM, Université PSL, 75005 Paris, France; Directors’ Research European Molecular Biology Laboratory, 69117 Heidelberg, Germany; Department of Systems Biology, Centro Nacional de Biotecnología (CNB), CSIC, Madrid, Spain; Red Sea Research Center, King Abdullah University of Science and Technology (KAUST), Thuwal 23955-6900, Saudi Arabia; Computational Bioscience Research Center (CBRC), King Abdullah University of Science and Technology (KAUST), Thuwal 23955-6900, Saudi Arabia; Departament de Genètica i Microbiologia, Facultat de Biociències, Universitat Autònoma de Barcelona, 08193 Bellaterra, Spain

**Keywords:** bacterial isolates, heterotrophic bacteria, prokaryotic communities, bathypelagic ocean, 16S iTAGs amplicons, sinking particles

## Abstract

Traditional culture techniques usually retrieve only a small fraction of the environmental marine microbial diversity, which mainly belong to the so-called rare biosphere. However, this paradigm has not been fully tested at a broad scale, especially in the deep ocean. Here, we examined the fraction of heterotrophic bacterial communities in photic and deep ocean layers that could be recovered by culture-dependent techniques at a large scale. We compared 16S rRNA gene sequences from a collection of 2003 cultured isolates of heterotrophic marine bacteria with global 16S rRNA metabarcoding datasets (16S TAGs) covering surface, mesopelagic and bathypelagic ocean samples that included 16 of the 22 samples used for isolation. These global datasets represent 60,322 unique 16S amplicon sequence variants (ASVs). Our results reveal a significantly higher proportion of isolates identical to ASVs in deeper ocean layers reaching up to a 28% of the 16S TAGs of the bathypelagic microbial communities, which included the isolation of 3 of the top 10 most abundant 16S ASVs in the global bathypelagic ocean, related to the genera *Sulfitobacter, Halomonas and Erythrobacter*. These cultured isolates contributed differently to the prokaryotic communities across different plankton size fractions, recruiting between 38% in the free-living size fraction (0.2-0.8 μm) and up to 45% in the largest plankton size fraction (20-200 μm) in the bathypelagic ocean. Our findings support the hypothesis that sinking particles in the bathypelagic realm act as resource-rich habitats, suitable for the growth of heterotrophic bacteria with a copiotroph lifestyle that can be cultured, and that these cultivable bacteria can also thrive as free-living bacteria.

## Introduction

Microbial ecology has long been limited by the fact that only a small fraction of the natural bacterial communities can be cultivated, a phenomenon that has traditionally been called “the great plate count anomaly” (Staley and Konopka, 1985). The recovered proportion of cells using selective media and standard plating techniques when compared to microscopy counts by direct staining is thought to represent only among 0.001-1 % of the community (Razumov, 1932; Kogure *et al.*, 1979; Staley and Konopka, 1985; Amann *et al.*, 1995). This phenomenon led to the known paradigm that “less than 1% of the microbial cells can be cultured” (Jannasch, 1958; Kuznetsov, 1975; Staley and Konopka, 1985; Eguchi and Ishida, 1990). However, few studies have attempted to isolate bacteria including mesopelagic and bathypelagic samples at a large scale, and therefore the long-standing observation that traditional culture techniques only retrieve a small fraction of the microbial diversity in marine environments still needs to be properly tested in the pelagic deep ocean.

Various seminal studies comparing culture-dependent and culture-independent techniques in marine ecosystems, revealed, in most cases, that there is a very small overlap between isolates and environmental sequences (Floyd *et al.*, 2005; Zeng *et al.*, 2012; Lekunberri *et al.*, 2014), and that most bacterial strains growing under laboratory conditions belong to the rare biosphere (Shade *et al.*, 2012; Crespo *et al.*, 2016). Nevertheless, these studies are scarce, and mostly focused on the photic ocean (0-200 m) (Eilers *et al.*, 2000; Zeng *et al.*, 2012; Lekunberri *et al.*, 2014; Crespo *et al.*, 2016), or in specific deep-ocean ecosystems such as hydrothermal vents (Harmsen *et al.*, 1997; Huber *et al.*, 2002; Hirayama *et al.*, 2007), hypersaline deep-sea basins (Sass *et al.*, 2001), or deep-sea sediments (Gutierrez *et al.*, 2015; Chen *et al.*, 2016), leaving the mesopelagic and bathypelagic waters less explored.

Besides, some meta-analysis studies have analyzed both the abundance and the diversity of the prokaryotic community that can be isolated in different ecosystems, including marine environments (Lloyd *et al.*, 2018; Martiny, 2019; Steen *et al.*, 2019). However, in these studies, the deep ocean was poorly examined, and the authors used different genetic thresholds and methods for calculating which percentage of the prokaryotic diversity could be isolated generating contrasting results. Most importantly, they did not compare isolates and 16S amplicon TAGs from exactly the same samples. Therefore, in such comparisons it is missing the detection of those members of the prokaryotic community that can be isolated using traditional culture techniques and the description of how abundant they are in those same ocean samples.

Diversity of bathypelagic ocean microbial communities across the tropical and temperate oceans described the most abundant operational taxonomic units (OTUs) (Salazar *et al.*, 2016) and interestingly, some of these OTUs affiliated with well-known heterotrophic bacterial genera known to be easily retrievable in culture, including *Alteromonas, Alcanivorax* and *Halomonas* (Salazar *et al.*, 2016; Kai *et al.*, 2017; Sanz-Sáez *et al.*, 2020). Some of these deep-ocean marine bacteria, such as those from within the genus *Alteromonas,* are copiotrophs (López-Pérez *et al.*, 2012) and given that the bathypelagic realm is fed by sinking particles (Herndl and Reinthaler, 2013), which are likely resource-rich habitats for microbes (Bochdansky *et al.*, 2016; Boeuf *et al.*, 2019). We hypothesize that their copiotrophic lifestyle could be a relevant survival strategy for some of the abundant bacteria inhabiting the deep ocean. Altogether, we postulate that i) higher proportions of the bacteria dwelling in the deep ocean would be retrieved under laboratory conditions and that ii) particles in the deep ocean are hotspots of copiotrophic bacteria that may be more easily isolated in culture than the free-living ones or than those present at the surface.

## Results

We examined the fraction of heterotrophic microbial communities that could be retrieved by isolation from both the photic and the deep ocean and explore how abundant are cultured bacteria in different plankton size fractions. To that end, we combined results from a large collection of heterotrophic cultured bacteria (MARINHET_v2), covering a wide range of oceanographic regions and depths, including the photic, the mesopelagic and the bathypelagic ocean (Sanz-Sáez *et al.*, 2020), with culture-independent results including flow cytometry measurements and 16S metabarcoding datasets obtained from simultaneous samples as the ones used for isolation from global oceanographic expeditions, *Tara* Oceans 2009-2013 (including *Tara* Oceans Polar Circle samples) (Sunagawa *et al.*, 2020) and Malaspina Circumnavigation Expedition 2010 (Ruiz-González *et al.*, 2019). Details regarding all samples used in this study can be found in **Figure 1**.

**Figure 1.**
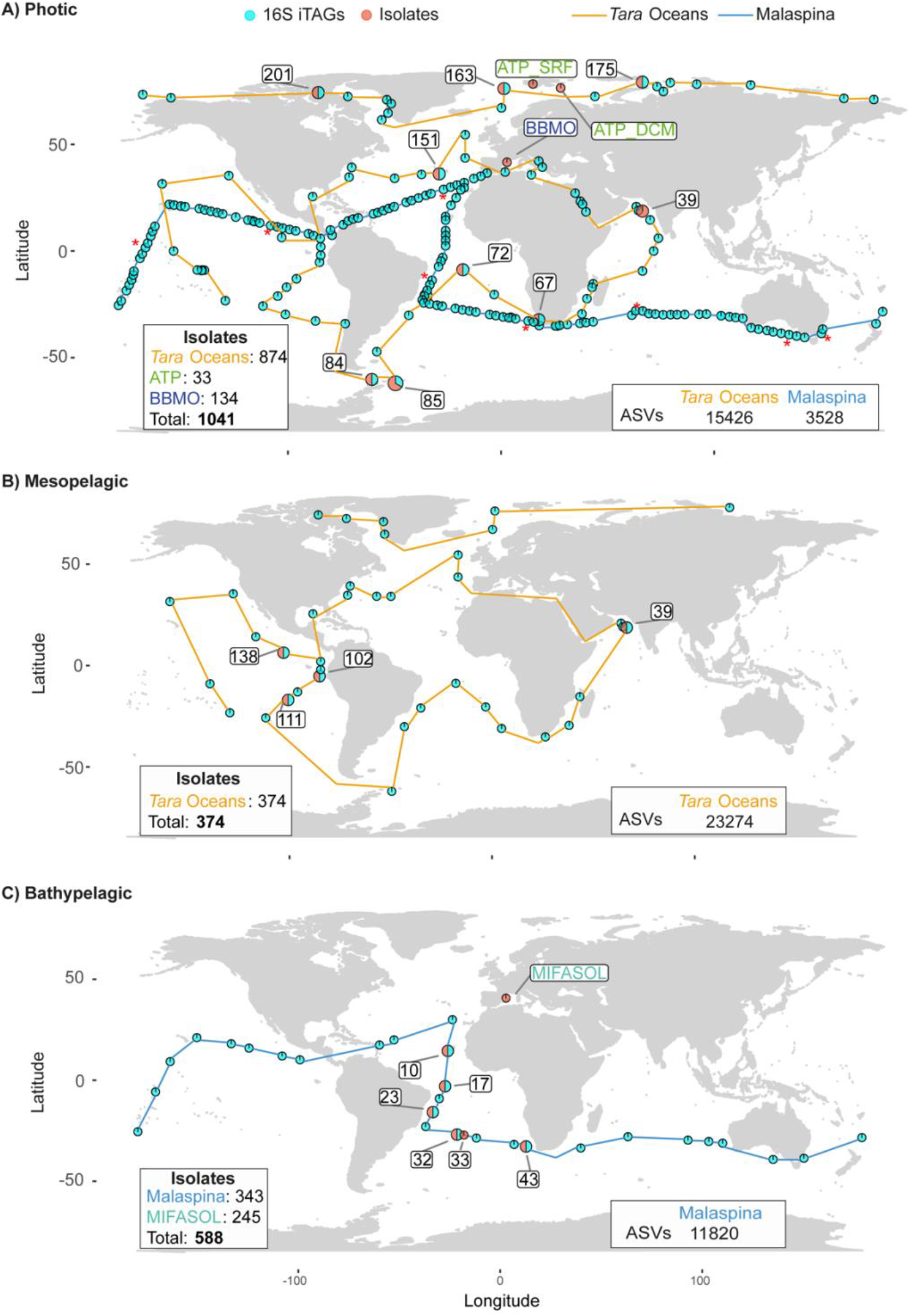
World map showing the distribution of the samples used in this study per ocean layer. **(A)** Photic. Labeled samples correspond to those stations where isolates were obtained from: *Tara* Oceans (39, 67, 72, 76, 84, 85, 151, 163, 175, 201), Blanes Bay Microbial Observatory (BBMO), and ATP Arctic cruise (ATP_SRF, ATP_DCM). Stations with red asterisks correspond to amplicon 16S iTAGs from eight vertical profiles with five different microbial size fractions collected from the Malaspina Expedition. **(B)** Mesopelagic. Labeled samples correspond to those stations where isolates were obtained from the *Tara* Oceans (39, 102, 111, 138). **(C)** Bathypelagic. Labeled samples correspond to those stations where isolates were obtained from: Malaspina (10, 17, 23, 32, 43) and MIFASOL. Circles connected with a blue line show the distribution of the samples obtained from the Malaspina Expedition, while circles connected with an orange line show those from the *Tara* Oceans. Each pie chart shows the presence or absence of samples from the different datasets: orange, isolates and light-blue, 16S iTAGs.

### Testing the “great plate count anomaly” in different oceanic regions and depths

We calculated the percentage of isolated bacterial cells for 10 photic-layer, 3 mesopelagic and 7 bathypelagic stations where plate colony counts (cfu/ml) and flow cytometry values - as a measure of total prokaryotic abundance/concentration- (cells/ml) were available (**Figure 2**). For this comparative analysis, flow cytometry counts included only the abundance of all heterotrophic prokaryotic and excluded photosynthetic *Cyanobacteria* since these taxa were not targeted by the media or incubation conditions selected. Considering that, we detected a higher percentage of recovery in the mesopelagic and bathypelagic samples (1.3 %) compared to the photic layer (0.3 %), although the differences were not significant (P > 0.05). The percentage of cultivability of heterotrophic bacterial cells ranged from 0.01 to 1.3 % in photic-layer samples, and from 0.9 to 2.5 % in mesopelagic samples, while percentages for bathypelagic samples varied between 0.08 % and 3.5 %, indicating a higher success in isolation in some samples from the deeper layers of the ocean (**Figure 2**).

**Figure 2.**
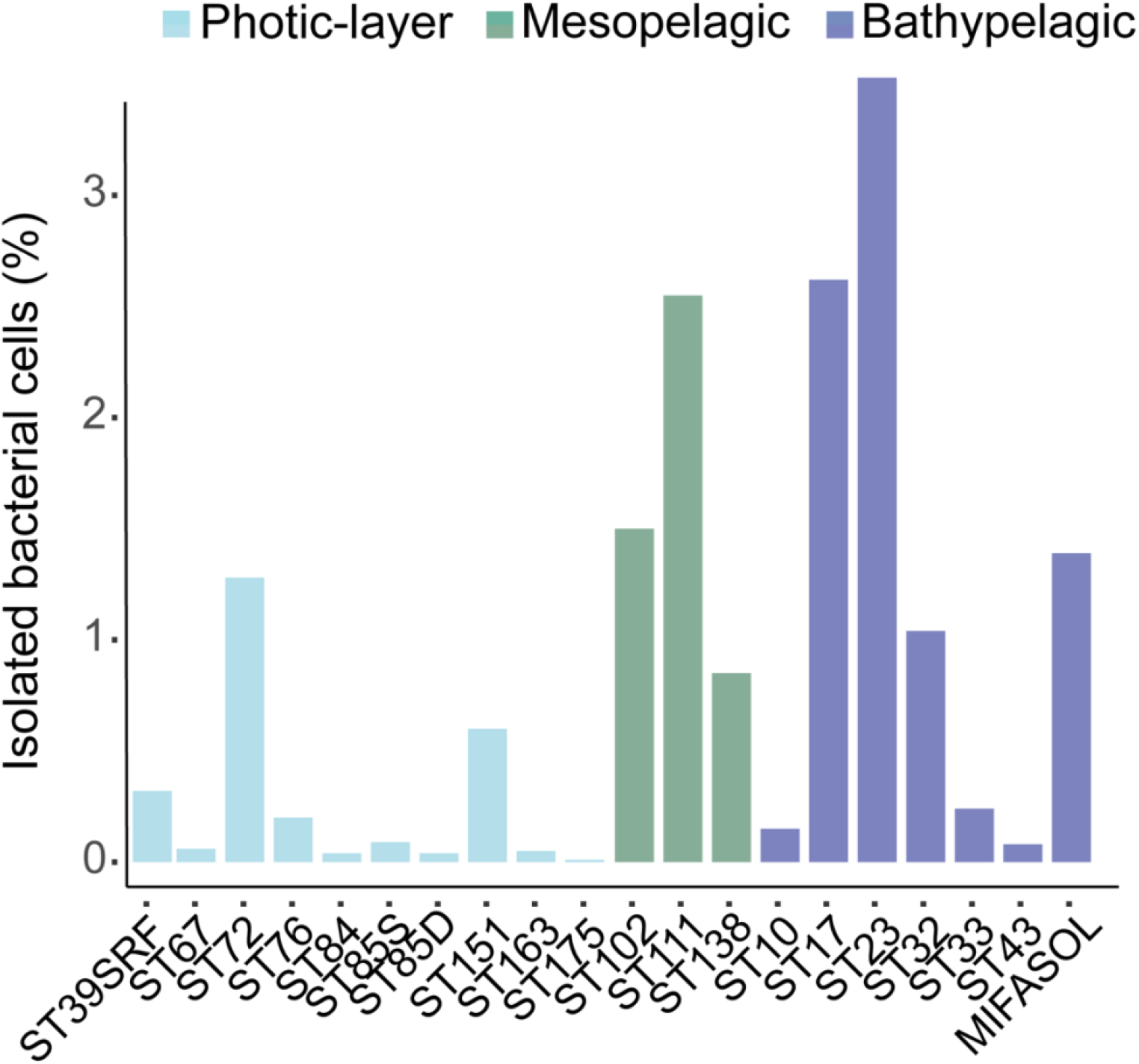
Testing the great plate count anomaly in different oceanic regions and ocean layers. For each station, heterotrophic prokaryotic abundances were estimated using both, traditional culture techniques (cfu/ml) and flow cytometry (cells/ml). The percentages represent the fraction of heterotrophic prokaryote abundance that could be retrieved by culturing in photic, mesopelagic and bathypelagic stations. Color indicates the depth of the sample: cyan, photic-layer; turquoise, mesopelagic; and dark-blue, bathypelagic. No significative differences were found between layers (ANOVA, P-value > 0.05).

### Contribution of culturable bacteria to total prokaryotic diversity in different ocean layers

A total of 2003 heterotrophic isolates were compared with two global metabarcoding datasets (16S TAGs) amplified with the same primer set (515F-Y-926R) (Parada et al. 2016) with a total of 38,700 ASVs from *Tara* Oceans (15,426 *Tara Ocenas* Surface and 23,274 *Tara Oceans* Mesopelagic) and 15,348 ASVs from the Malaspina Expedition (3,528 Malaspina Surface and 11,820 Malaspina Bathypelagic) (**Supplementary Table S1**, **Figure 1**). We determined the mean percentage of ASVs (diversity) as well as the mean percentage of 16S TAGs (abundance) that were 100% identical to our MARINHET_v2 culture collection isolates. These comparisons were done at three different levels: (i) comparing each 16S TAGs dataset from *Tara* Oceans and Malaspina Expedition with all the isolate sequences, (ii) comparing separately the photic and aphotic 16S TAGs datasets with only photic or aphotic zone isolate sequences, and (iii) comparing the 16S TAGs and the rRNA gene sequences of the isolates retrieved from exactly the same stations. A summary of the results of these three levels of comparisons, obtained from the rarefied ASVs-abundance table, is shown in **Supplementary Table S2**.

The highest average number of ASVs that were 100% identical to the isolates was observed in the Malaspina Surface dataset (4.5%), followed by the Malaspina Bathypelagic (2.4%), *Tara* Oceans Surface (2.3%) and *Tara* Oceans Mesopelagic datasets (1.7%) (**Figure 3A**). Even though these percentages do not seem to vary greatly between datasets, significant differences were found among them (Kruskal-Wallis, P-value < 0.01, **Figure 3A**). Similarly, when the comparisons of the datasets were performed separately for the photic and aphotic zone isolates, the highest mean % of ASVs equal to isolates was found in Malaspina Surface (2.7% photic, 4.1% aphotic) and the lowest percentages were found in *Tara* Oceans Mesopelagic (0.9% photic, 1.1% aphotic) (**Supplementary Table S2).** Otherwise, if we consider the abundances of these ASVs and we look at the percentage of reads (16S TAGs) identical to the isolates, we saw a significant increase in the deep ocean (**Figure 3B**). Thus, around 1.6-4.9 % of the 16S TAGs were 100% identical to our isolates in the photic ocean, this value increased up to 8.5% in the mesopelagic ocean, and increased even more, up to 27.9%, in the bathypelagic ocean. In this case, the differences between datasets were also statistically significant (Kruskal-Wallis test, P-value < 0.01, **Figure 3B**). Differences were also found when comparing all the isolates together or comparing separately photic and aphotic zone isolates (**Figure 3C-D** and **Supplementary Table S2**). The metabarcoding 16S TAGs dataset of Malaspina Bathypelagic samples integrates free-living (0.2-0.8 μm) and particle-attached (0.8 – 20 μm) microbial communities. However, if we use only the data from the free-living fraction to be fair with the comparisons with the free-living bacteria analyzed from the photic and mesopelagic samples, despite being slightly different size fractions (0.2-0.8 μm in Malaspina Bathypelagic vs. 0.2-3 μm in *Tara* Oceans Mesopelagic), we still observed this trait of higher proportions, with 22.9% of the 16S TAGs 100% identical to isolates in the bathypelagic samples (**Supplementary Figure S1**).

**Figure 3.**
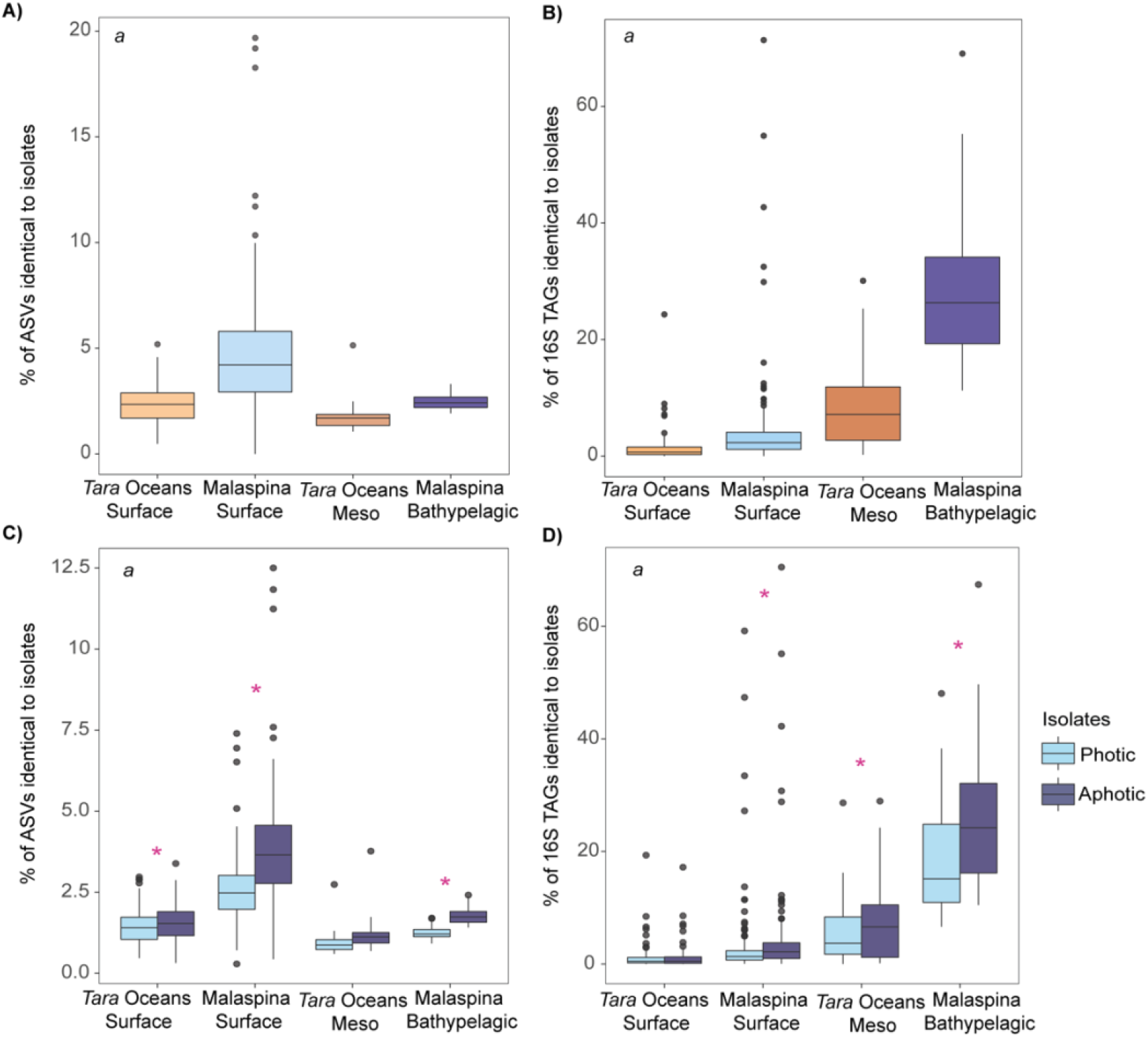
**(A) Proportion of isolates that matched at 100% identity to ASVs. (B) Proportion of 16S TAGs reads matching isolates at 100 % similarity.** Values are extracted from the mean abundance of reads or ASVs in each dataset from rarified ASV-abundance tables. Outliers are indicated with grey circles. If significant differences are found between all datasets it is indicated inside boxplots with an italic *a* (Kruskal-Wallis, P-value < 2.2 e-16). **(C) Proportion of isolates that are 100% similar to the ASVs comparing photic and aphotic zone isolates separately. (D) Proportion of 16S TAGs reads matching isolates at 100% similarity when comparing photic and aphotic zone isolates separately.** Significative differences between datasets (P-value < 0.01) are indicated by an italic *a* in the top left corner, while significative differences (P-value < 0.01) within a dataset are indicated by pink asterisks.

A fraction of the isolates did not match any ASV regardless of the dataset inspected (**Supplementary Table S3**). Indeed, approximately 11% of our heterotrophic isolates did not match any of the *Tara* Oceans ASVs, whereas this number increased up to 18% in the Malaspina Bathypelagic ASVs and up to 28% in the surface samples from Malaspina (**Supplementary Figure S2**).

Some interesting taxonomic differences at the family level were observed between the isolates that were identical to ASVs and those that did not match any ASV (**Supplementary Figure S3).** We found some families that were identified in the *Tara* Oceans Surface and Mesopelagic metabarcoding datasets but not in the Malaspina Expedition Surface or Bathypelagic metabarcoding datasets, such as *Tistrellaceae, Nitrincolaceae or Colwelliaceae.* In contrast, *Kangiellaceae* was found in of both Malaspina Expedition datasets but not in *Tara* Oceans samples. Some families included isolates that did not match any ASVs in any of the 16S metabarcoding datasets, such as *Dermabacteraceae*, *Balneolaceae* or *Psychromonadaceae* **(Supplementary Figure S3)**. These results indicate that despite the large sequencing effort used to obtain amplicon 16S TAGs in *Tara* Oceans (~ 5×10^5^ samples average reads) and in the Malaspina Expedition samples (surface samples ~ 5.1×10^4^ average reads, bathypelagic ~9.7×10^5^ average reads), there was still a proportion of the isolates, which may belong to the extremely rare biosphere, that could not be detected by this sequencing technique.

### Bacterial isolates among the most abundant ASVs of the deep ocean

To test whether isolates belonged to the abundant or rare biosphere, rank abundance plots were done for each of the *Tara* Oceans and Malaspina Expedition metabarcoding datasets, with the ASVs mean abundances extracted from rarefied ASVs-abundance tables (**Figure 4**). The rank abundances were plotted based on either the global comparison of all isolates (**Figure 4)** or by comparing photic and aphotic zone isolates separately against photic 16S TAGs datasets, or photic and aphotic zone isolates separately versus mesopelagic and bathypelagic 16S TAGs datasets (**Supplementary Figure S4**) giving similar results in all of them. Each rank abundance plot showed a similar pattern of few abundant ASVs (relative abundance > 1%), relatively few mid-abundant ASVs (<1% and >0.01%) and a long tail of rare or low-abundant ASVs (relative abundance <0.01%). We colored the ASVs that were 100% similar to at least one isolate to test for differences between depths (**Figure 4**). In photic layers, we did not have any isolate within the abundant taxa of the *Tara* Oceans Surface dataset (**Figure 4A**), whereas only one belonging to the abundant biosphere was found in the Malaspina Surface dataset, taxonomically related to *Sulfitobacter* with 1.7% of the total reads (**Figure 4B).** The rest of the isolates in these two large-scale photic datasets appeared in the mid-abundant biosphere (38 in Malaspina and three in *Tara Oceans*) or in the rare biosphere (54 in Malaspina and 151 in *Tara Oceans*). In the mesopelagic layer (*Tara Oceans* Mesopelagic) we only found one ASV identical to isolates classified into the abundant biosphere (associated with *Alteromonas* with 1.1% of the reads), although more isolates were identical to ASV with medium abundances (87 ASVs) (**Figure 4C**). Interestingly, the most abundant bathypelagic ocean taxon according to the 16S TAGs matched at 100% identity with one of our isolates. This organism was related to *Sulfitobacter* and represented 4.6% TAGs reads. In total, seven ASV identical to isolates belonged to the abundant biosphere in Malaspina Bathypelagic, affiliating with the genera *Sulfitobacter*, *Halomonas*, *Erythrobacter, Alteromonas and Sphingobium,* and the first three were included into the top 10 most abundant ASV detected. These, together with a relatively large proportion of isolates that matched organisms of the mid-abundant biosphere (78 ASVs) organisms, and those matching rare biosphere organisms (52), recruited 28% of the environmental 16S TAGs from the temperate and tropical global bathypelagic oceans (**Figure 4D**). Thus, abundant ASVs could be retrieved by culture-dependent techniques, especially in the bathypelagic layer.

**Figure 4.**
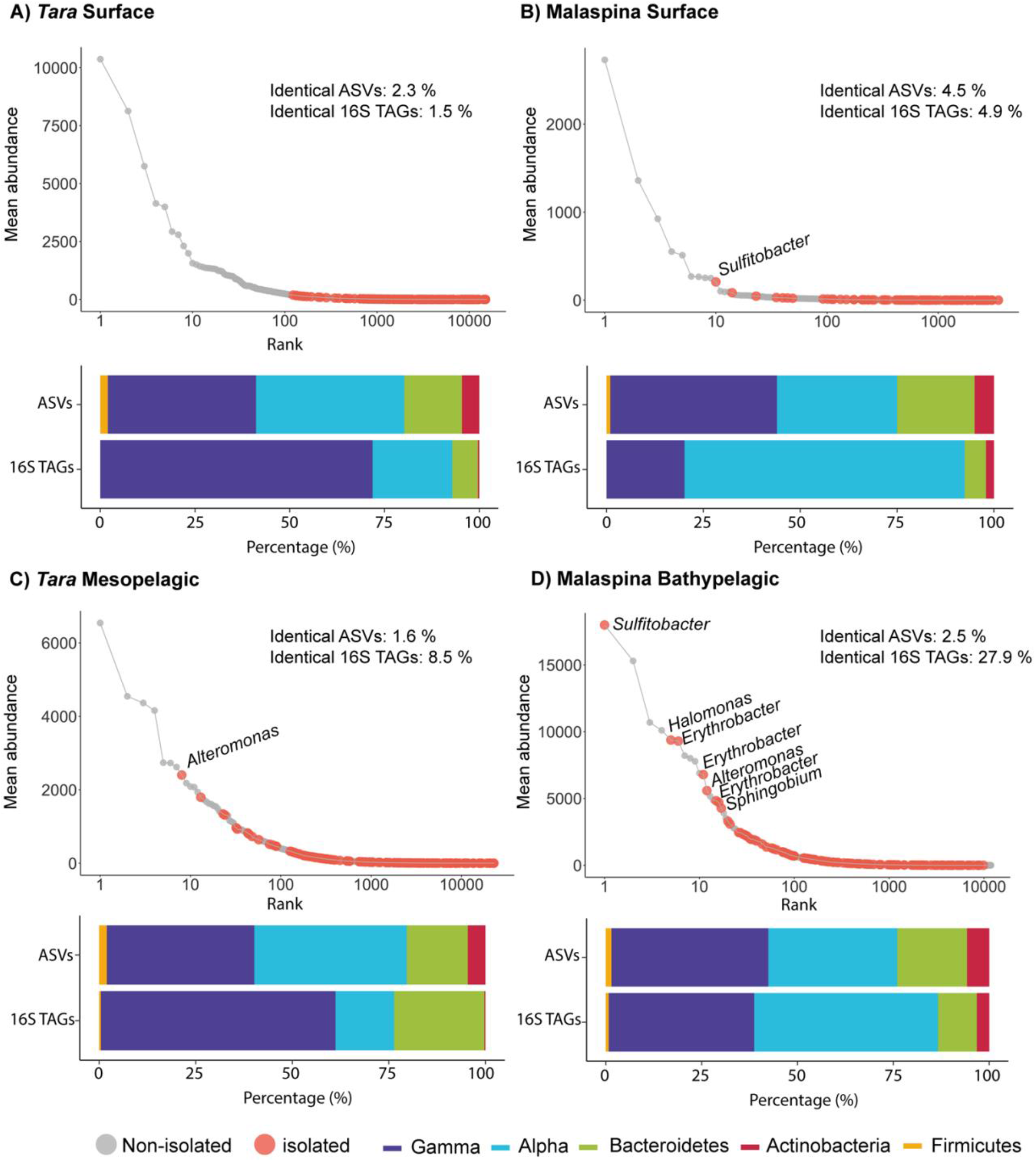
Rank plots showing the identified isolates that matched at 100% identity to 16S iTAGs in different datasets. **(A)** *Tara* Oceans Surface **(B)** Malaspina Surface. **(C)** *Tara* Oceans Mesopelagic **(D)** Malaspina Bathypelagic. Coloured dots indicates ASVs matched by isolates: grey, isolates that did not matched any ASVs; orange, isolates with 100% identity to ASVs. Taxonomic affiliation is indicated for the abundant (>1 % abundance) isolates that matches with ASVs. The histograms describe the proportions of the isolates that marched with ASVs or the proportion of 16S TAGs (reads) at the phylum/class level: dark-blue, *Gammaproteobacteria;* light-blue, *Alphaproteobacteria;* green, *Bacteroidetes;* red, *Actinobacteria;* and orange, *Firmicutes.*

Altogether, we found that the ASVs that were 100% identical to our isolates affiliated mostly with classes *Alphaproteobacteria* (average 39% ASVs) and *Gammaproteobacteria* (average 47.7% ASVs) followed by phyla *Bacteroidetes* (average 11.5%), *Actinobacteria* (average 1.5%) and *Firmicutes* (average 0.3%) (**Figure 4** and **Supplementary Table S4**). We noticed that despite finding relatively similar proportions of isolates belonging to *Alphaproteobacteria* and *Gammaproteobacteria* in all ocean layers, the proportion of reads within these isolates identical to ASVs of these classes differed between the *Tara* Oceans and Malaspina Expedition datasets regardless of sampling depth. In *Tara* Oceans samples, *Gammaproteobacteria* dominated (72%-61%), while *Alphaproteobacteria* dominated in the Malaspina datasets (72%-48%) (**Figure 4**). These differences could be explained because relatively different ocean regions, latitudes and seasons were sampled by each expedition, in general more coastal regions were sampled in *Tara* Oceans, but also Malaspina Expedition arrived to deeper layers (up to 4000 m depth) compared to *Tara* Oceans.

### Increase in isolated sequences from larger plankton size fractions

We also aimed to elucidate whether a higher proportion of isolates could be retrieved from bacteria developing on particles in the deep ocean representing hotspots of copiotrophic bacteria, with the hypotheses that particle-attached (PA) bacteria would be more easily isolated in culture than the free-living (FL) ones or than those present at the surface. We observed a significant increase (Wilcoxon test, P < 0.01) of isolates 100% identical to ASVs and of reads in the PA communities (0.8 – 20 μm, average 32.6%, sd:13.3%) versus the FL fraction (0.2 – 0.8 μm, average 22.99%, sd: 10.2%) (**Supplementary Figure S1**) in the Malaspina Bathypelagic samples. However, the PA community analyzed in the Malaspina Bathypelagic dataset corresponded to particles of many different sizes from 0.8 to 20 μm plankton size fraction. Therefore, to explore the effect of the particle size range on the percentage of ASVs recovery, we also compared our isolates with samples from five different plankton size fractions (0.2-0.8 μm - considered FL; or PA in sizes: 0.8-3.0 μm, 3.0-5.0 μm, 5.0-20 μm, and 20-200 μm) in eight vertical profiles from the Malaspina Expedition.

Our analysis revealed that the isolates were present across all size fractions and depths, yet the proportions varied (**Figure 5** and **Supplementary Figure S5**). First, when looking at the differences between layers (surface, DCM, mesopelagic and bathypelagic), we confirmed again the higher average diversity (% of ASVs) and abundance (% of TAGs) of isolated bacteria in the bathypelagic. The 16.3% of ASVs 100% identical to isolates from the bathypelagic (**Figure 5A**) recruited an average of 40% of TAGs (**Figure 5B**). Among surface, DCM and mesopelagic samples, a similar number of isolates were detected (~ 8% average) and also similar proportions of identical TAGs were identified (~ 20% average) (**Figure 5**). Interestingly, the DCM was the layer with the smaller number of ASVs identical to isolates (~ 6% average) and TAGs recruited (~17.3% on average). Given the notable proportion of ASVs identical to isolates in the bathypelagic samples, statistically significant differences were found between this deeper layer and the other depths in the different size fractions (Kruskall-Wallis test, P-values <0.01), but not between the surface, DCM or mesopelagic samples (**Supplementary Figures S5-S6**).

**Figure 5.**
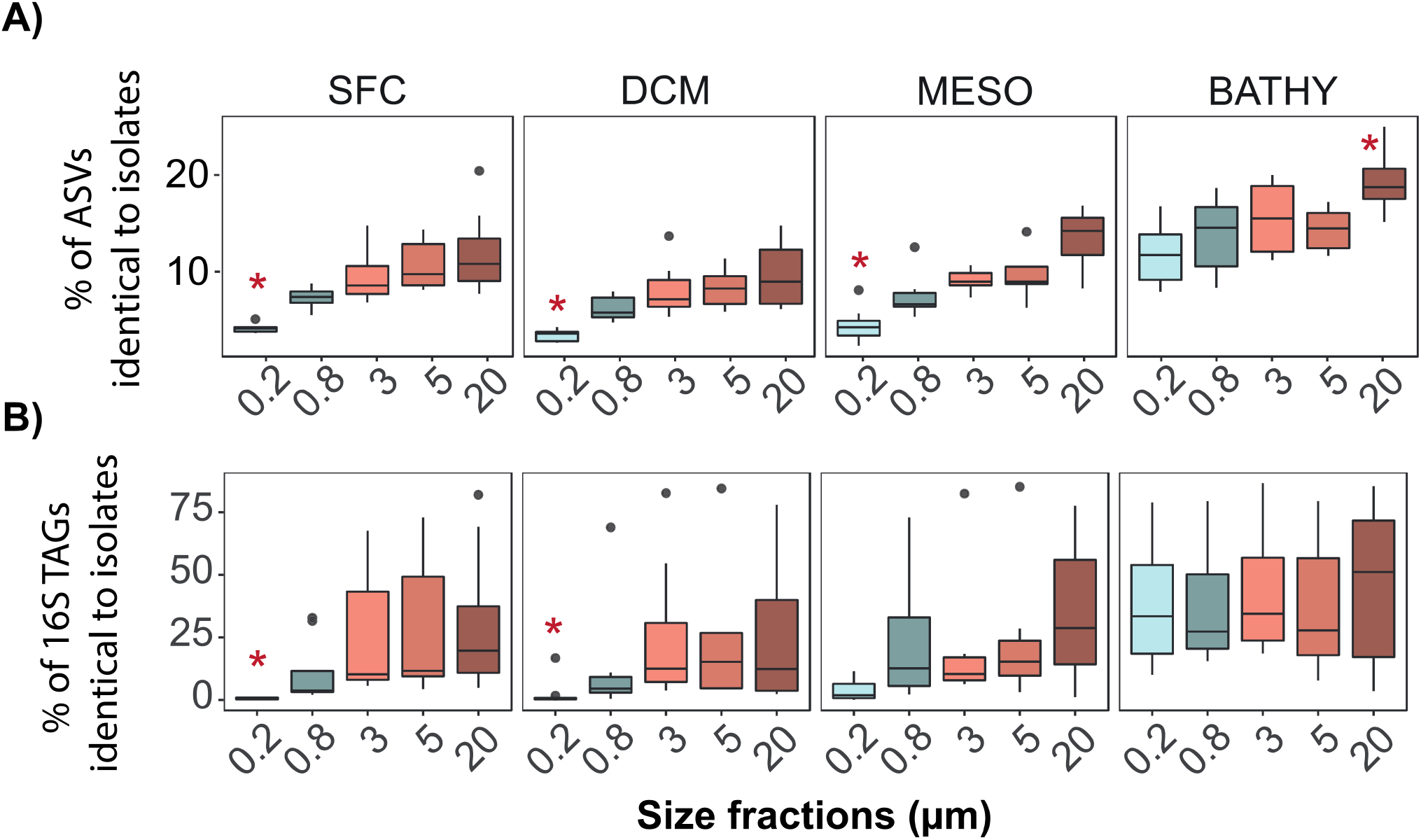
Boxplots comparing five different plankton size fractions in the Malaspina Expedition vertical profiles: **(A)** percent of isolates identical to ASVs per size fraction and depth, and **(B)** percentage of reads (16S iTAGs) that were 100% identical to at least one isolate. SFC: surface, DCM: deep chlorophyll maximum, MESO: mesopelagic, and BATHY: bathypelagic. 0.2 μm: free-living bacteria (0.2-0.8 μm), 0.8 μm: bacteria attached to small particles (0.8-3 μm), and 3.0-20 μm: bacteria attached to larger particles. Significant differences between size fractions in each layer are indicated by red asterisks (Kruskal-Wallis, P.value <0.05

Additionally, the proportion of sequences identical to isolates across different plankton size fractions, revealed that our isolates were prevalent in the larger plankton size fractions, those associated with particles (≥3.0 μm) (**Figure 5**). In samples from the photic zone (surface and DCM) higher mean abundances of ASVs and TAGs (10% and 26% on average, respectively) were recovered by isolation in the larger size fractions (3.0 μm, 5.0 μm and 20 μm) but not in the FL bacterial communities (0.2 μm, 2.9% average ASVs, and 1.5% average TAGs) or in the smallest particles (0.8 μm, 6% average ASVs, and 12% average TAGs) (**Supplementary Table S5** and **S6**). In the mesopelagic samples, the number of ASVs identical to isolates (7%) in the 0.8 μm size fraction recruited up to 23% of the TAGs, but not in the FL fraction (0.2 μm), which still presented lower values (4% of reads). Finally, in the bathypelagic samples, all plankton size fractions displayed similar percentages of ASVs identical to isolates, including the FL bacteria (average 16%), and uniform proportions of TAGs (40% on average). In addition, the highest values were found from the largest particles (>20 μm, 21% of ASVs and 45% of the TAGs).

## Discussion

Culturing studies are still fundamental for microbial ecologists to fully understand the ecology of microorganisms in marine ecosystems and test hypothesis that with sequencing techniques alone cannot be fully answered. In this study, combination of culture-dependent and culture-independent techniques have allowed us to test the great plate count anomaly paradigm in the deep ocean including mesopelagic and bathypelagic zones, or to test the hypothesis weather bathypelagic bacteria and those bacteria associated to particles are more prone to be isolated in pure culture.

The precise meaning of the traditional paradigm that only 1% of microbes are culturable has a difficult interpretation as discussed by Martiny (2019). Our comparative analyses between the number of colonies retrieved in pure culture and the cell abundances calculated with flow cytometry provided information about the proportion of cells (abundance) than could be recovered by isolation using a specific set of culture and incubation conditions. Our results for mesopelagic and bathypelagic samples reflected a certain degree of variability, with a mean percentage of cultivability between 1.5% and 3.7%. This variability has also been highlighted in studies that examined culture-dependent surveys covering a wide variety of environments: lakes, seawater, soils or sediments, and also human host-associated communities (Lloyd *et al.*, 2018; Steen *et al.*, 2019). However, none of the studies reviewed by these authors included open ocean samples deeper than 200 m depth. We can confirm that this variance can also be observed across the mesopelagic and the bathypelagic oceans, and that more than 1% of the cells and up to approximately 4% of them (**Figure 2**) could be cultivated in some samples from the deep ocean. Therefore, the traditional paradigm that less than 1% of the microbial cells can be retrieved in culture should be not literally interpreted. However, related but different questions are: which fraction of the microbial community diversity can be cultured?, how representative of the global community are the isolated bacteria?, and are the isolates abundant or rare members of the microbial community? To get these answers we compared the isolates 16S rRNA sequences at the maximal resolution possible (at 100% identity) with metabarcoding16S TAGs from the same ocean samples.

The rank abundance curves of microbial communities are composed by some high-abundant and moderately abundant taxa but many low abundant taxa, the so called rare biosphere (Sogin *et al.*, 2006; Pedrós-Alió, 2012). We confirmed this structure for both expeditions datasets and different depth levels in the ocean, in agreement with previous studies (Pommier et al. 2010, Sunagawa *et al.*, 2015; Crespo *et al.*, 2016; Salazar *et al.*, 2016). The common idea that cultivation mainly captures members of the rare biosphere seems to be the rule for bacteria in the photic layer, as the majority of the isolates from *Tara* Oceans and Malaspina (an average 81% isolates) recruited less than 1% of the total 16S rRNA sequences at 100% identity in datasets from the photic zone (**Figure 3**). These results were also observed in a single station in the northwestern Mediterranean Sea, in which 24% of the isolates from surface seawater were found in the amplicon data (454 tags), yet all belonged to rare taxa representing globally less than 1% of the total reads (Crespo *et al.*, 2016).

More interestingly, we found that 2.4% of the isolates from the bathypelagic ocean recruited up to 28% of the reads of the total microbial community in the deep ocean realm identified by amplicon 16S TAGs (**Figure 3B**) reflecting that we are capturing some of the abundant taxa of the deep ocean. Our isolation strategy focused only on the heterotrophic marine bacteria and we are aware that the photosynthetic bacterial community in the surface samples is not captured with our isolation strategy nor are the Archaea, which could represent an important fraction of the bathypelagic ocean (Salazar *et al.*, 2016). We tested what occurred when removing *Cyanobacteria* and Archaea TAGs from the datasets, and we obtained a similar trend with a slight increase and identifying a similar average of the amplicon 16S TAGs reads identical to isolates in the surface, mesopelagic and bathypelagic datasets (**Supplementary Table S7**).

On the other hand, the 16S metabarcoding datasets used in this study are obtained with a PCR-dependent method, which could have influenced the results obtained given that isolated organisms usually have a higher rRNA operon copy number, and thus, overestimating the abundance of TAGs recruited by the ASV identical to our isolates. This bias can be corrected by dividing our results by 3.5, which is the median rRNA operon copy number of the isolated genera in this study (**Supplementary Table S8**). Then, the proportion of TAGs recruited by ASV 100% identical to isolates is reduced in *Tara Oceans* Surface from 1.6% to 0.5%, in Malaspina Surface from 4.8% to 1.4%, in *Tara Oceans* Mesopelagic from 8.5% to 2.4% and in Malaspina Bathypelagic from 27.9% to 8 %. Even though the abundance of the isolated community diminishes, still higher proportions are found in the deeper layers and within those organisms we found some of the top10 ASV detected among our culture collection, such as the most abundant ASV in the bathypelagic ocean affiliating to *Sulfitobacter* genera.

Our study also highlights that the recruitment of isolates was higher in the particle attached fraction of all layers, and especially in the largest size fractions of the bathypelagic ocean (**Figure 5B**). It is already well know that marine microbial communities attached to particles are very different from the free-living ones both, taxonomically (Acinas *et al.*, 1999; Eloe *et al.*, 2011; Crespo et al. 2013, Salazar *et al.*, 2015; Mestre *et al.*, 2018), and at the functional level (Smith *et al.*, 2013; López-Pérez *et al.*, 2016; Acinas *et al.*, 2021).

A previous study that analyzed the same Malaspina size fractionated vertical profiles dataset (Mestre *et al.*, 2018) determined the importance of sinking particles as promoters of vertical connectivity in the ocean microbiome and observed that bacteria associated with particles in the surface belonging to the rare taxa became dominant in the deep ocean. Our results show that marine heterotrophic cultured bacteria, despite being rare taxa in the surface ocean, are capable of growing in these hotspot particles and to become abundant in the deep ocean. Thus, we noticed in the bathypelagic samples similar proportions of recruited reads by our isolates in all size fractions, suggesting that our heterotrophic isolates represent well the bathypelagic realm in all plankton size fractions, varying from 38% of the reads in the free-living smallest size fraction (0.2-3 μm) up to 45% in the largest particles (20-200 μm). The observed similarities between size fractions in the bathypelagic ocean could be due to the presence of isolates with dual lifestyles, i.e., the same isolate is capable of living in large particles and as a free-living bacteria, as has been reported for some marine *Flavobacteria* (González *et al.*, 2008). Additionally, a recent study on the globally active bathypelagic microbiome that combined 16S rRNA and 16S rDNA metabarcoding revealed the dominance of prokaryotes with dual lifestyles (Sebastián *et al.*, 2021). Furthermore, the isolates that were abundant (>1% relative abundance) in both free-living and particle-attached bacterial communities in the bathypelagic samples were also present in the largest size fractions of the surface layers, but appeared in the mid-abundant and rare biosphere of the free-living fraction (**Supplementary Figure S7** and **Table S9**). Surface bacteria could thus act as a seed bank for bathypelagic communities, a hypothesis that has also been proposed in other studies (Mestre et al. 2016; Gibbons *et al.*, 2013; Sebastián *et al.*, 2017; Ruiz-González *et al.*, 2020). While our isolates likely belonged to the part of the bacterial community that prefers a particle-attached lifestyle, our results support that they could live attached to particles in surface waters and sink with the particles to deeper layers where they would develop and finally become abundant members of the planktonic community **(Figure 6).** We are therefore retrieving in pure culture isolates adapted to live attached to particles with a copiotroph lifestyle, where nutrients are abundant and still can survive as free-living bacteria in the bathypelagic deep ocean.

**Figure 6.**
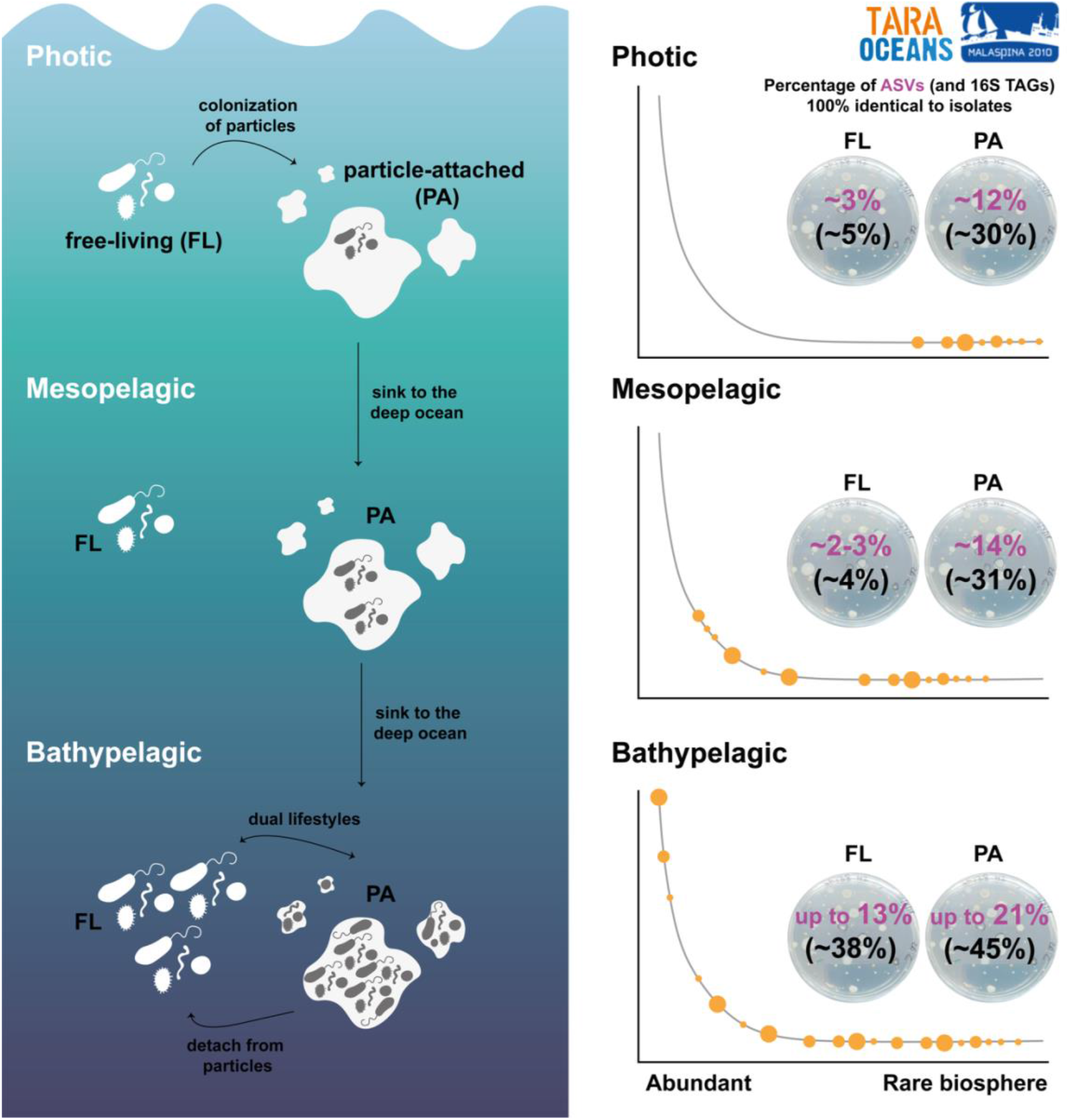
Conceptual representation of the heterotrophic culturable bacteria along the ocean water column. Free-living bacteria from photic ocean, which present a small fraction of heterotrophic culturable bacteria and mainly belonging to the rare biosphere, can attached to particles, where this culturable fraction is higher. These particles serve as a hotspot for growing and sink into the deep ocean where heterotrophic culturable bacteria become more abundant specially in larger particles. Once in the deep ocean, mainly in the bathypelagic, bacteria can detach from particles and present a dual-lifestyle. In the bathypelagic, some of the most abundant bacteria are culturable and they are present both in the free-living and particle attached fraction.

Particles are resource-rich habitats for microbes (Bochdansky *et al.*, 2016) where a copiotroph lifestyle could be the rule. Our findings support this idea given the fact that we found a higher proportion of isolates identical to the ASVs detected in the particle-attached communities, especially in the bathypelagic communities. Also genomic comparisons between cultured isolates and uncultured genomes retrieved by single amplified genomes (SAGs) from marine environments showed that the genomes of the cultured bacteria had larger sizes, with a predominant copiotrophic lifestyle (Swan *et al.*, 2013) which could have favored us the recovery of the particle-attached bacteria using the culture and incubation conditions selected. Some of the most abundant ASVs identical to isolates detected in the bathypelagic ocean affiliating to *Sulfitobacter, Halomonas* and *Alteromonas* display higher growth rates (>4 day^-1^) than other heterotrophic bacteria (Fecskeová *et al.*, 2021), which may also favored the isolation in pure culture. However, our MARINHET_v2 culture collection still misses most of the free-living bacteria adapted to oligotrophic conditions in the photic and the mesopelagic layers. Future isolation efforts using dilution-to-extinction (Rappé *et al.*, 2002; Selje *et al.*, 2005), microdroplets or cultivation chips (Ingham *et al.*, 2007), methods combining cell sorting with isotopic labeling (Thrash, 2021), or the use of metagenomic and metatranscriptomic data to predict the metabolic requirements of certain bacterial groups and adapt culture media accordingly (Gutleben *et al.*, 2017) may help us bring to the laboratory some other abundant biogeochemically key taxa of the free-living microbial communities that remain mostly uncultured. Nevertheless, the genome sequencing of these deep ocean abundant taxa in combination with physiological experiments under different scenarios of temperature, pression, nutrients or particles association would enhance our understanding of key taxa of the deep ocean.

## Conclusions

This is the first comparative large-scale study of culture-dependent and culture-independent techniques from the same water samples from different oceanographic regions and depths (including the meso- and the bathypelagic). Our findings confirmed the traditional paradigm that isolation techniques retrieve members of the rare biosphere in the photic ocean although the percentage of recovery can increase up to 3% of the cells in the bathypelagic ocean. Most importantly, this study revealed that ecologically relevant abundant bacteria can be retrieved by isolation, especially in the bathypelagic ocean, across all plankton size fractions, but mainly in those associated with larger particles where we found values representing up to 45% of the TAGs and representing some of the most abundant members of the community.

## Material and Methods

### Isolates culture collection database

Water samples from a total of 14 photic-layer stations and 11 deep-ocean stations (including 4 from the mesopelagic in oxygen minimum zone (OMZ) regions and 7 from the bathypelagic) were collected from global oceanographic expeditions including the Malaspina Expedition (Duarte, 2015) and *Tara* Oceans (*Tara* Oceans 2009 and *Tara* Oceans Polar Circle 2013)(Sunagawa *et al.*, 2020). Additionally, we used seawater samples collected for isolation in other cruises, such as ATP09 in the Arctic Ocean (Lara *et al.*, 2013), MIFASOL in the NW Mediterranean, as well as from the Blanes Bay Microbial Observatory (BBMO, http://www.icm.csic.es/bio/projects/icmicrobis/bbmo), covering a wide latitudinal range, with different oceanographic regions. Seawater samples collected for isolation were prefiltered through 200 μm and 20 μm mesh in succession in order to keep free-living bacteria but also the prokaryotic community attached to particles < 20 μm. Further details regarding sample collection in all these cruises have been previously described in Sanz-Sáez et al. (2020). Geographical coordinates of the stations, sampled depth, *in situ* temperature, number of isolates sequenced, total prokaryote cell abundances, and colony forming units (cfu) per mL are listed in Supplementary **Table S10.** Prokaryote cell abundance was determined using flow cytometry (in a BectonDickinson FACSCalibur) of SYBR Green I stained samples (Gasol and Morán, 2016).

Our culturing strategy focused on retrieving heterotrophic marine bacteria that could easily grow under laboratory conditions (nutrient rich media, standard oxygen concentrations and atmospheric pressure) (**Supplementary Table S11**). Therefore, we used nutrient rich media including Zobell agar, Marine Agar 2216 and modified Marine Agar, where disodium phosphate was autoclaved separated from the rest of the media and added as a separate solution before solidification (Zobell, 1941; Sanz-Sáez *et al.*, 2020). Further details regarding isolation of bacterial strains has been already described in (Sanz-Sáez *et al.*, 2020).

A total of 2003 bacterial isolates (MARINHET_v2 culture collection) were randomly selected for DNA amplification and partial sequencing of their 16S rRNA gene (more details in (Sanz-Sáez *et al.*, 2020)). Similar number of isolates were selected from photic layers (1041; average: 70.6 isolates per station) and from the deep ocean (962; average: 67.6 isolates per station).

### Metabarcoding 16S rRNA datasets

We used 54,048 amplicon sequence variants (ASVs), obtained from several datasets: Malaspina Surface (124 samples)(Ruiz-González *et al.*, 2019), Malaspina Bathypelagic (41 samples, average depth: 3731 m ± 495; standard deviation)(Salazar *et al.*, 2016), *Tara* Surface (80 samples) and *Tara* Mesopelagic (39 samples) (this study and Ibarbalz *et al.*, 2019) **(Figure 1),** and an additional 6,274 ASVs derived from eight vertical profiles generated in the Malaspina Expedition that included 5 different size fractions for four depths corresponding to surface (3 m), the depth of the deep chlorophyll maximum (DCM, 48-150 m), mesopelagic (250-670 m), and bathypelagic waters (3105-4000 m) **(Figure 1A)**(Mestre *et al.*, 2018). These samples covered most tropical and temperate ocean regions but also some polar oceanic regions (*Tara* Oceans Polar Circle expedition **Figure 1A,B)**. The ASVs were compared to the MARINHET_v2 culture collection that contains 2003 isolates (including those already present in the first version already published in (Sanz-Sáez *et al.*, 2020) and an extra set of 442 isolates) retrieved from different oceanographic regions and depths, including photic, mesopelagic and bathypelagic waters **(Figure 1).**

All samples were collected with 20 L Niskin bottles and prefiltered through 200 μm and 20 μm mesh in succession. Volumes filtered and filters used for collecting prokaryotic DNA for analyses of the bacterial community using 16S amplicon Illumina TAGs are specified in **Table 1** for each cruise and type of sampling. Filters were then flash-frozen in liquid nitrogen and stored at −80 °C until DNA extraction.

**Table 1.** Volumes and filters used for collecting prokaryotic DNA. Fractions analyzed in each dataset it is also indicated.

|  | Malaspina Surface (3m) | Malaspina Bathypelagic | Malaspina vertical-profiles* | *Tara* Oceans and *Tara* Oceans Polar Circle |
| --- | --- | --- | --- | --- |
| Volume (L) | 6-15 | 120 | 10 | 100 |
| Prefilters (µm) | 200/20 | 200/20 | 200/20 | 200/20 |
| Filters (µm) | 3.0/0.2 | 0.8/0.2 | 5.0/3.0/0.8/0.2 | 3/0.2 or 1.6/0.2 |
| Fractions analyzed | 0.2 – 3 | 0.2 – 0.8<br>0.8 – 20 | 0.2 – 0.8<br>0.8 – 3.0<br>3.0 – 5.0<br>5.0 – 20<br>20 – 200 | 0.2 – 1.6<br>Or<br>0.2 – 3.0 |

\* Surface, deep chlorophyll maximum (DCM), mesopelagic, and bathypelagic samples DNA extraction and sequencing for metabarcoding datasets

The samples for 16S metabarcoding sequencing were extracted with a phenol-chloroform protocol (as described in Massana *et al.*, 1997; Salazar *et al.*, 2016; Alberti *et al.*, 2017). Prokaryotic barcodes for each of the datasets were generated by amplifying the V4 and V5 hypervariable regions of the 16S rRNA gene using primers 515F-Y (5’-GTGYCAGCMGCCGCGGTAA-3’) and 926R (5’-CCGYCAATTYMTTTRAGTTT-3’) described in Parada et al. (2016). The Malaspina bathypelagic DNA samples originally from Salazar et al. (2016) were re-sequenced again using 515F-Y-926R primers (Parada *et al.*, 2016) to be comparable with the rest of the analyzed 16S metabarcoding datasets in this study. Sequencing for *Tara* Oceans and *Tara* Oceans Polar Circle and Malaspina Bathypelagic datasets was performed at Genoscope using an Illumina MiSeq platform (iTAGs) with the 2×250 bp paired-end approach. The Malaspina Surface and Malaspina vertical-size fraction profiles datasets were sequenced at the Research and Testing Laboratory facility (https://rtlgenomics.com) also with Illumina MiSeq platform and the 2×250 bp paired-end approach.

### Metabarcoding sequence data processing

All 16S rRNA amplicons were processed *de novo* through the bioinformatic pipeline described in the GitHub repository (https://github.com/SushiLab/Amplicon_Recipes) regardless they were previously analyzed and published. Briefly, pair-end reads were merged at a minimum 90% of identity alignment, and those with ≤ 1 expected errors were selected (quality filtering). Primer matching was performed with CUTADAPT v.1.9.1. Dereplication, definition of zero-radius OTUs or amplicon sequence variants (ASVs) were performed with USEARCH v.10.0.240 (Edgar, 2010) using UNOISE3 algorithm. ASVs were taxonomically annotated against the SILVA database v.132 (2018) with the lowest common ancestor (LCA) approach. Finally, we built ASV-abundance tables. Non-prokaryotic ASVs (eukaryotes, chloroplasts and mitochondria) were removed, whereas singletons (ASVs appearing only once) were maintained. For some specific analyses, Cyanobacteria or Archaea reads were also removed since our isolation protocol was not adequate for culturing these microorganisms. Computing analyses were run at the MARBITS bioinformatics platform at the *Institut* de *Ciències del Mar* and at the Euler scientific compute cluster of the ETH Zürich University.

This procedure was applied individually for the: (i) 41 Malaspina Bathypelagic samples, (ii) 124 Malaspina Surface samples, (iii) 119 *Tara* Oceans and *Tara* Oceans Polar Circle Surface and Mesopelagic samples, and (iv) 155 samples from the Malaspina vertical-size fractions profiles dataset. Hence, four different and new ASV-abundance tables were obtained after applying the above pipeline (Malaspina datasets were re-preprocessed from the original publications of the samples (Salazar *et al.*, 2016; Mestre *et al.*, 2018; Ruiz-González *et al.*, 2019)). Each ASV table was randomly sampled down to lowest sampling effort using the function *rrarefy.perm* with 1000 permutations from the R package *EcolUtils* (Salazar, 2018). A summary of the total number of reads per dataset, sample with the lowest number of reads and total ASVs 100% before and after rarefication is described in Supplementary **Table S1**.

### Comparison between 16S amplicon TAGs and cultured isolates

The primers used to obtain the 16S rRNA genes of the isolates were different from the ones used to obtain the 16S rRNA gene TAGs, although both amplified the V4 and V5 hypervariable region of the 16S rRNA gene. Therefore, comparisons between isolates and 16S TAGs were performed by selecting this common region (**Supplementary Figure S8**).

All isolate sequences were compared to the ASVs at 100% similarity in order to have the strictest comparison possible. Comparisons were done by running global alignments using the *usearch_global* option from the USEARCH v10.0.240 (Edgar, 2010). The results were filtered by coverage of the alignment at 100% (i.e. all the ASVs sequences must align without any gaps in the sequence with the partial sequences of the isolates). We are aware that these comparisons sometimes result in more than one sequence hit per isolate. Nevertheless, all datasets contained similar proportions of isolates with more than one sequence match and the % of reads recruited by these extra hits was minor (approximately 1.2% in each dataset), making comparisons between datasets possible (**Supplementary Figure S2**).

### Data analysis

All data analyses were done with the R Statistical Software (R core team, 2017) using v.3.4.3 and the following packages: *vegan* (Oksanen *et al.*, 2018), *ape* (Paradis *et al.*, 2004), *EcolUtils* (Salazar, 2018), *stats* (R core team, 2017), *tidyverse* (Wickham, 2019). For each dataset we calculated the mean abundance and relative abundance of ASVs across samples in order to rank the organisms detected. Moreover, we calculated the mean percentage of 16S TAGs reads (how much the isolates represent the bacterial community in terms of abundance?) and the ASVs (how much the isolates represent the bacterial community in terms of diversity or richness?) of the bacterial community that matched at 100% similarity with the 16S rRNA gene sequences of the strains isolated by traditional culture techniques. These percentages were calculated from the rarefied ASV-abundance tables. In order to test the significance of the differences between the proportion of isolates that matched the amplicon reads of each dataset, we used non-parametric Kruskall-Wallis followed by post hoc pairwise Wilcox test. To assess significance, the statistical analyses were set to an alpha value of 0.05.

We used the 16S TAGs dataset from the Malaspina vertical-size fractionated vertical profiles to investigate whether our isolates were enriched in the free-living microbial fraction (0.2-0.8 μm), or in the particle-attached community, considering that this last category can be divided into four different size-fractions (0.8-3 μm, 3-5 μm, 5-20 μm and 20-200 μm). Hence, we also calculated the percentage of reads or 16S TAGs, and ASVs that matched at 100% similarity with our isolates. The differences between size fractions were also tested with the non-parametric Kruskall-Wallis test followed by the post hoc pairwise Wilcox test to determine the differences between pairs of datasets. Again, the significance was set at an alpha value of 0.05.

### Nucleotide accession numbers

The isolates sequences are deposited in GenBank under accession numbers MH731309 - MH732621 and MK658870-MK659428. Amplicon 16S rRNA iTAGs from the different datasets used are available in the European Nucleotide Archive (ENA). Those from the Malaspina Surface dataset are available under accession number PRJEB25224, those from the Malaspina Bathypelagic dataset under PRJEB45011, those from Malaspina size fractions under PRJEB27154 and those from *Tara* Oceans and *Tara* Oceans Polar Circle under accession numbers PRJEB36282, PRJEB36283 and PRJEB36439.

## Acknowledgments

We thank our fellow scientists and the crew and chief scientists of cruises ATP, MIFASOL and Malaspina and all scientist involved in the monthly sampling at the BBMO. We thank the people and sponsors who participated in the *Tara* Oceans Expedition 2009–2013 (http://oceans.taraexpeditions.org) for sampling collection. The project Malaspina 2010 Expedition (ref. CSD2008–00077) was funded by the Spanish Ministry of Economy and Competitiveness, Science and Innovation through the Consolider-Ingenio program. Other funding support were from Projects Arctic Tipping Points (ATP, contract #226248), in the FP7 program of the European Union, and DOREMI (CTM2012–34294) from the Spanish Ministry of Economy and Competitiveness, which allowed the collection of samples from the Arctic and NW Mediterranean Sea, respectively. Research, including laboratory experiments and analyses, was mainly funded by grant MAGGY, Plan Nacional I + D + I 2017 (CTM2017–87736-R), Polar EcoGen, Plan Nacional I+D+I 2020 (PID2020-116489RB-I00), and the Swiss National Science Foundation (SNSF) through project grant 205321_184955. Further funding was obtained from the King Abdullah University of Science and Technology (KAUST) under subaward agreement OSR#3362 and project MIAU-S3 (ref. RTI2018–101025-B-I00) from the Spanish Ministry of Economy and Competitiveness. This work benefited from the institutional support of the ‘Severo Ochoa Centre of Excellence’ accreditation (CEX2019-000928-S).

## Notes

### Competing Interest Statement

The authors have declared no competing interest.

## References

Acinas, S.G., Antón, J., and Rodríguez-Valera, F. (1999) Diversity of free-living and attached bacteria in offshore Western Mediterranean waters as depicted by analysis of genes encoding 16S rRNA. Appl Environ Microbiol 65: 514–522.

Acinas, S.G., Sánchez, P., Salazar, G., Cornejo-Castillo, F.M., Sebastián, M., Logares, R., et al. (2021) Deep ocean metagenomes provide insight into the metabolic architecture of bathypelagic microbial communities. Commun Biol 4: 604.

Alberti, A., Poulain, J., Engelen, S., Labadie, K., Romac, S., Ferrera, I., et al. (2017) Viral to metazoan marine plankton nucleotide sequences from the Tara Oceans expedition. Sci data 4: 170093.

Amann, R.I., Ludwig, W., and Schleifer, K.H. (1995) Phylogenetic identification and in situ detection of individual microbial cells without cultivation. Microbiol Rev 59: 143–169.

Bochdansky, A.B., Clouse, M.A., and Herndl, G.J. (2016) Dragon kings of the deep sea: marine particles deviate markedly from the common number-size spectrum. Sci Rep 6: 22633.

Boeuf, D., Edwards, B.R., Eppley, J.M., Hu, S.K., Poff, K.E., Romano, A.E., et al. (2019) Biological composition and microbial dynamics of sinking particulate organic matter at abyssal depths in the oligotrophic open ocean. Proc Natl Acad Sci U S A 116: 11824–11832.

Chen, P., Zhang, L., Guo, X., Dai, X., Liu, L., Xi, L., et al. (2016) Diversity, biogeography, and biodegradation potential of Actinobacteria in the deep-sea dediments along the Southwest Indian ridge. Front Microbiol 7: 1340.

Crespo, B.G., Wallhead, P.J., Logares, R., and Pedrós-alió, C. (2016) Probing the rare biosphere of the North-West Mediterranean Sea: an experiment with high sequencing effort. PLoS One 11: e0159195.

Duarte, C.M. (2015) Seafaring in the 21St Century: The Malaspina 2010 Circumnavigation Expedition. Limnol Oceanogr Bull 24: 11–14.

Edgar, R.C. (2010) Search and clustering orders of magnitude faster than BLAST. Bioinformatics 26: 2460–2461.

Eguchi, M. and Ishida, Y. (1990) Oligotrophic properties of heterotrophic bacteria and in situ heterotrophic activity in pelagic seawaters. FEMS Microbiol Ecol 73: 23–30.

Eilers, H., Pernthaler, J., Glöckner, F.O., and Amann, R. (2000) Culturability and in situ abundance of pelagic Bacteria from the North Sea. Appl Environ Microbiol 66: 3044–3051.

Eloe, E.A., Shulse, C.N., Fadrosh, D.W., Williamson, S.J., Allen, E.E., and Bartlett, D.H. (2011) Compositional differences in particle-associated and free-living microbial assemblages from an extreme deep-ocean environment. Environ Microbiol Rep 3: 449–458.

Fecskeová, L.K., Piwosz, K., Šantic, D., Šestanovic, S., Tomaš, A.V., Hanusová, M., et al. (2021) Lineage-specific growth curves document large differences in response of individual groups of marine bacteria to the top-down and bottom-up controls. mSystems 6:.

Floyd, M.M., Tang, J., Kane, M., and Emerson, D. (2005) Captured diversity in a culture collection: case study of the geographic and habitat distributions of environmental isolates held at the american type culture collection. Appl Environ Microbiol 71: 2813–2823.

Gasol, J.M. and Morán, X.A.G. (2016) Flow cytometric determination of microbial abundances and its use to obtain indices of community structure and relative activity. In, McGenity,T.J., Timmis,K.N., and Nogales,B. (eds), drocarbon and Lipid Microbiology Protocols. Springer Protocols Handbooks. Springer, pp. 159–187.

Gibbons, S.M., Caporaso, J.G., Pirrung, M., Field, D., Knight, R., and Gilbert, J.A. (2013) Evidence for a persistent microbial seed bank throughout the global ocean. Proc Natl Acad Sci U S A 110: 4651–4655.

González, J.M., Fernández-Gómez, B., Fernàndez-Guerra, A., Gómez-Consarnau, L., Sánchez, O., Coll-Lladó, M., et al. (2008) Genome analysis of the proteorhodopsin-containing marine bacterium Polaribacter sp. MED152 (Flavobacteria). Proc Natl Acad Sci U S A 105: 8724–8729.

Gutierrez, T., Biddle, J.F., Teske, A., and Aitken, M.D. (2015) Cultivation-dependent and cultivation-independent characterization of hydrocarbon-degrading bacteria in Guaymas Basin sediments. Front Microbiol 6: 1–12.

Gutleben, J., De Mares, M.C., Dirk Van Elsas, J., Smidt, H., Overmann, J., and Sipkema, D. (2017) The multi-omics promise in context: from sequence to microbial isolate. Crit Rev Microbiol 44: 212–219.

Harmsen, H.J.M., Prieur, D., and Jeanthon, C. (1997) Distribution of microorganisms in deep-sea hydrothermal vent chimneys investigated by whole-cell hybridization and enrichment culture of thermophilic subpopulations. Appl Environ Microbiol 63: 2876–2883.

Herndl, G.J. and Reinthaler, T. (2013) Microbial control of the dark end of the biological pump. Nat Geosci 6: 718–724.

Hirayama, H., Sunamura, M., Takai, K., Nunoura, T., Noguchi, T., Oida, H., et al. (2007) Culture-dependent and -independent characterization of microbial communities associated with a shallow submarine hydrothermal system occurring within a coral reef off Taketomi Island, Japan. Appl Environ Microbiol 73: 7642–7656.

Huber, J.A., Butterfield, D.A., and Baross, J.A. (2002) Temporal changes in archaeal diversity and chemistry in a mid-ocean ridge subseafloor habitat. Appl Environ Microbiol 68: 1585–1594.

Ingham, C.J., Sprenkels, A., Bomer, J., Molenaar, D., van den Berg, A., van Hylckama Vlieg, J.E.T., and de Vos, W.M. (2007) The micro-Petri dish, a million-well growth chip for the culture and high-throughput screening of microorganisms. Proc Natl Acad Sci 104: 18217–18222.

Jannasch, H.W. (1958) Studies on planktonic bacteria by means of a direct membrane filter method. J gen Microbiol 18: 609–620.

Kai, W., Peisheng, Y., Rui, M., Wenwen, J., and Zongze, S. (2017) Diversity of culturable bacteria in deep-sea water from the South Atlantic Ocean. Bioengineered 8: 572–584.

Kogure, K., Simidu, U., and Taga, N. (1979) A tentative direct microscopic method for counting living marine bacteria. Can J Microbiol 25: 415–420.

Kuznetsov, S.I. (1975) The microflora of lakes and its geochemical activity, Oppenheimer,C.H. (ed) Austin and London: University of Texas Press 1976.

Lara, E., Arrieta, J.M., Garcia-Zarandona, I., Boras, J.A., Duarte, C.M., Agustí, S., et al. (2013) Experimental evaluation of the warming effect on viral, bacterial and protistan communities in two contrasting Arctic systems. Aquat Microb Ecol 70: 17–32.

Lekunberri, I., Gasol, J.M., Acinas, S.G., Gómez-Consarnau, L., Crespo, B.G., Casamayor, E.O., et al. (2014) The phylogenetic and ecological context of cultured and whole genome-sequenced planktonic bacteria from the coastal NW Mediterranean Sea. Syst Appl Microbiol 37: 216–228.

Lloyd, K.G., Steen, A.D., Ladau, J., Yin, J., and Crosby, L. (2018) Phylogenetically novel uncultured microbial cells dominate Earth microbiomes. mSystems 3: e00055–18.

López-Pérez, M., Gonzaga, A., Martin-Cuadrado, A.-B., Onyshchenko, O., Ghavidel, A., Ghai, R., and Rodriguez-Valera, F. (2012) Genomes of surface isolates of Alteromonas macleodii: the life of a widespread marine opportunistic copiotroph. Sci Rep 2: 696.

López-Pérez, M., Kimes, N.E., Haro-Moreno, J.M., and Rodriguez-Valera, F. (2016) Not all particles are equal: the selective enrichment of particle-associated Bacteria from the Mediterranean Sea. Front Microbiol 7: 996.

Martiny, A.C. (2019) High proportions of bacteria are culturable across major biomes. ISME J 13: 2125–2128.

Massana, R., Murray, A.E., Preston, C.M., and Delong, E.F. (1997) Vertical Distribution and Phylogenetic Characterization of Marine Planktonic Archaea in the Santa Barbara Channel. Appl Environ Microbiol 63: 50–56.

Mestre, M., Ruiz-González, C., Logares, R., Duarte, C.M., Gasol, J.M., and Sala, M.M. (2018) Sinking particles promote vertical connectivity in the ocean microbiome. Proc Natl Acad Sci U S A 115: E6799–E6807.

Oksanen, J., Blanchet, F.G., Friendly, M., Kindt, R., Legendre, P., McGlinn, D., et al. (2018) Vegan: community ecology package. R package version 2.5-3. https://CRAN.R-project.org/package=vegan.

Parada, A.E., Needham, D.M., and Fuhrman, J.A. (2016) Every base matters: Assessing small subunit rRNA primers for marine microbiomes with mock communities, time series and global field samples. Environ Microbiol 18: 1403–1414.

Paradis, E., Claude, J., and Strimmer, K. (2004) APE: analyses of phylogenetics and evolution in R language. Bioinformatics 20: 289–290.

Pedrós-Alió, C. (2012) The rare bacterial biosphere. Ann Rev Mar Sci 4: 449–466.

R core team (2017) R core team. A language and environment for statistical computing. R foundation for statistical computing, Vienna, Austria https://www.R-project.org/.

Rappé, M.S., Connon, S.A., Vergin, K.L., and Giovannoni, S.J. (2002) Cultivation of the ubiquitous SAR11 marine bacterioplankton clade. Nature 418: 630–633.

Razumov, A.S. (1932) The direct method of calculation of bacteria in water: comparison with the Koch method. Mikrobiologija 1: 131–146.

Ruiz-González, C., Mestre, M., Estrada, M., Sebastián, M., Salazar, G., Agustí, S., et al. (2020) Major imprint of surface plankton on deep ocean prokaryotic structure and activity. Mol Ecol 29: 1820–1838.

Ruiz-González, C., Logares, R., Sebastián, M., Mestre, M., Rodríguez-Martínez, R., Galí, M., et al. (2019) Higher contribution of globally rare bacterial taxa reflects environmental transitions across the surface ocean. Mol Ecol 28: 1930–1945.

Salazar, G. (2018) EcolUtils: Utilities for community ecology analysis. R package version 0.1. https://github.com/GuillemSalazar/EcolUtils.

Salazar, G., Cornejo-Castillo, F.M., Benítez-Barrios, V., Fraile-Nuez, E., Álvarez-Salgado, X.A., Duarte, C.M., et al. (2016) Global diversity and biogeography of deep-sea pelagic prokaryotes. ISME J 10: 596–608.

Salazar, G., Cornejo-Castillo, F.M., Borrull, E., Díez-Vives, C., Lara, E., Vaqué, D., et al. (2015) Particle-association lifestyle is a phylogenetically conserved trait in bathypelagic prokaryotes. Mol Ecol 24: 5692–5706.

Sanz-Sáez, I., Salazar, G., Sánchez, P., Lara, E., Royo-Llonch, M., Sà, E.L., et al. (2020) Diversity and distribution of marine heterotrophic bacteria from a large culture collection. BMC Microbiol 20: 207.

Sass, A.M.M., Sass, H., Coolen, M.J.J.L., Cypionka, H., and Overmann, J. (2001) Microbial communities in the chemocline of a hypersaline deep-sea basin (Urania basin, Mediterranean Sea). Appl Environ Microbiol 67: 5392–5402.

Sebastián, M., Auguet, J.-C., Restrepo-Ortiz, C.X., Sala, M.M., Marrasé, C., and Gasol, J.M. (2017) Deep ocean prokaryotic communities are remarkably malleable when facing long-term starvation. Environ Microbiol 20: 713–723.

Sebastián, M., Sánchez, P., Salazar, G., Álvarez-Salgado, X.A., Reche, I., Morán, X.A.G., et al. (2021) The quality of dissolved organic matter shapes the biogeography of the active bathypelagic microbiome. bioRxiv 2021.05.14.444136.

Selje, N., Brinkhoff, T., and Simon, M. (2005) Detection of abundant bacteria in the Weser estuary using culture-dependent and culture-independent approaches. Aquat Microb Ecol 39: 17–34.

Shade, A., Hogan, C.S., Klimowicz, A.K., Linske, M., McManus, P.S., and Handelsman, J. (2012) Culturing captures members of the soil rare biosphere. Environ Microbiol 14: 2247–2252.

Smith, M.W., Allen, L.Z., Allen, A.E., Herfort, L., Simon, H.M., and Nelson, C.E. (2013) Contrasting genomic properties of free-living and particle-attached microbial assemblages within a coastal ecosystem. Front Microbiol 4: 1–20.

Sogin, M.L., Morrison, H.G., Huber, J.A., Welch, D.M., Huse, S.M., Neal, P.R., et al. (2006) Microbial diversity in the deep sea and the underexplored “rare biosphere.” Proc Natl Acad Sci U S A 103: 12115–12120.

Staley, J.T. and Konopka, A. (1985) Measurement of in situ activities of nonphotosynthetic microorganisms in aquatic and terrestrial habitats. Annu Rev Microbiol 39: 321–346.

Steen, A.D., Crits-Christoph, A., Carini, P., Deangelis, K.M., Fierer, N., Lloyd, K.G., and Thrash, J.C. (2019) High proportions of bacteria and archaea across most biomes remain uncultured. ISME J 13: 3126–3130.

Sunagawa, S., Acinas, S.G., Bork, P., Bowler, C., Acinas, S.G., Babin, M., et al. (2020) Tara Oceans: towards global ocean ecosystems biology. Nat Rev Microbiol 18: 428–445.

Sunagawa, S., Coelho, L.P., Chaffron, S., Kultima, J.R., Labadie, K., Salazar, G., et al. (2015) Structure and function of the global ocean microbiome. Science (80-) 348: 1261359.

Swan, B.K., Tupper, B., Sczyrba, A., Lauro, F.M., Martinez-Garcia, M., González, J.M., et al. (2013) Prevalent genome streamlining and latitudinal divergence of planktonic bacteria in the surface ocean. Proc Natl Acad Sci U S A 110: 11463–11468.

Thrash, J.C. (2021) Towards culturing the microbe of your choice. Environ Microbiol Rep 13: 16–41.

Wickham, H. (2019) Welcome to the tidyverse. J Open Source Softw 4: 1686.

Zeng, Y., Zou, Y., Grebmeier, J.M., He, J., and Zheng, T. (2012) Culture-independent and -dependent methods to investigate the diversity of planktonic bacteria in the northern Bering Sea. Polar Biol 35: 117–129.

Zobell, C.E. (1941) Studies on marine bacteria. I. The requirements of heterotrophic aerobes. J Mar Res 4: 42–75.

