## Supplementary Material for "Abundant deep ocean heterotrophic bacteria are culturable"

### A significant fraction of the deep ocean heterotrophic bacteria are culturable

**This PDF file includes:**

Supplementary Methodology

Supplementary Figs. S1 to S9

Supplementary Tables S1 to S10

**Other Supplementary Material for this manuscript include the following:**

Excel file with Supplementary Tables S4

#### Supplementary Methodology

The results presented in this study are based on the detection of amplicon sequence variants (ASVs) compared to 16S rRNA isoltes sequences at 100% sequence similarity. However, during the analyses we also defined from metabarcoding 16S TAGs OTU at 97% with the UPARSE algorithm using USEARCH v.10.0.240 (Edgar, 2010). OTUs at 97% identity and zOTUs were taxonomically annotated against the SILVA database v.132 (2018) with the lowest common ancestor (LCA) approach. OTU-abundance tables were also defined for OTUs at 97% sequence similarity. OTUs at 97% sequence similarity were then compared to 16S rRNA isolates sequences at 97% sequence similarity. Some results about these comparisons can be found in Supplementary Table S1 and S2.

#### Supplementary Results

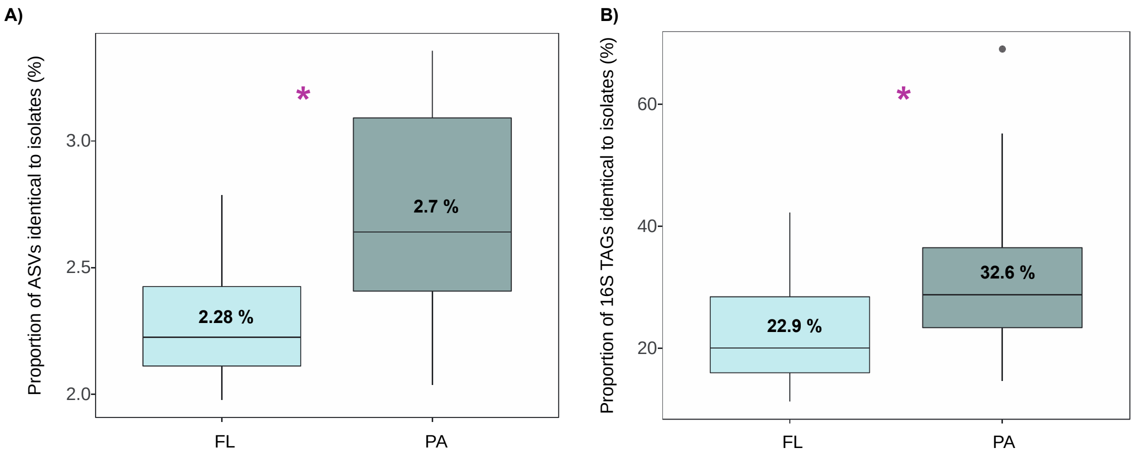

**Figure S1. Differences between size fractions in the Malaspina Bathypelagic datasets**. **(A)** Proportion of ASVs 100% identical to isolates. **(B)** Proportion of 16S TAGs (reads) identical to isolates. Comparisons were done separating the free-living (0.2-0.8 μm) and particle-attached (0.8-20 μm) communities. The average percentage is indicated inside the boxplots and significant differences between size fractions are indicated with pink asterisks (P-values < 0.05).

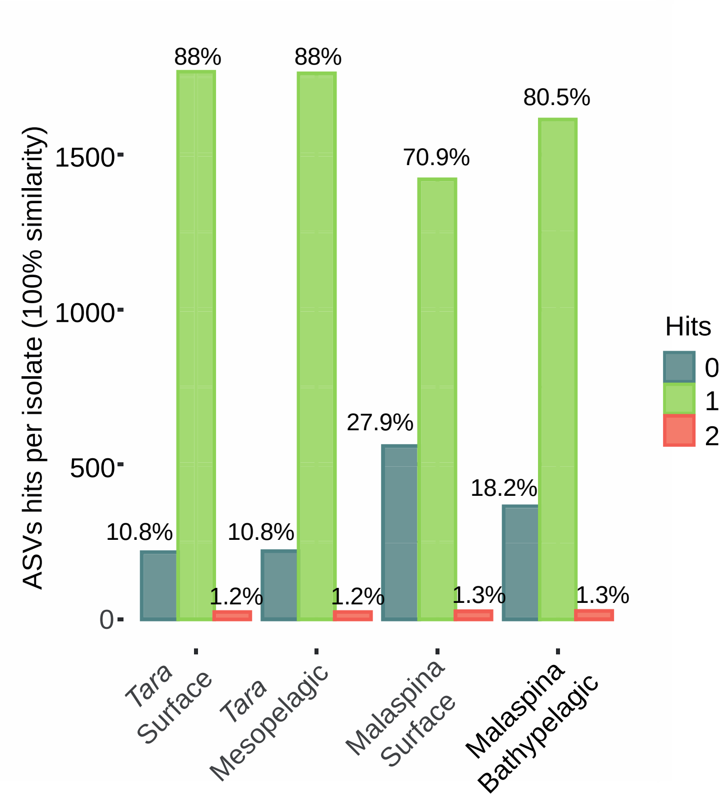

**Figure S2. Proportion of isolates identical to ASVs.** Number of isolates presenting no hits to any ASV (0 hits, dark-green), just hits to one single ASV (1 hit , pale green) or hits to 2 different ASVs (2 hits, orange). Comparisons done at 100 % sequence similarity using *usearch_global*. Percentages in each category from the total isolates are indicated above each bar.

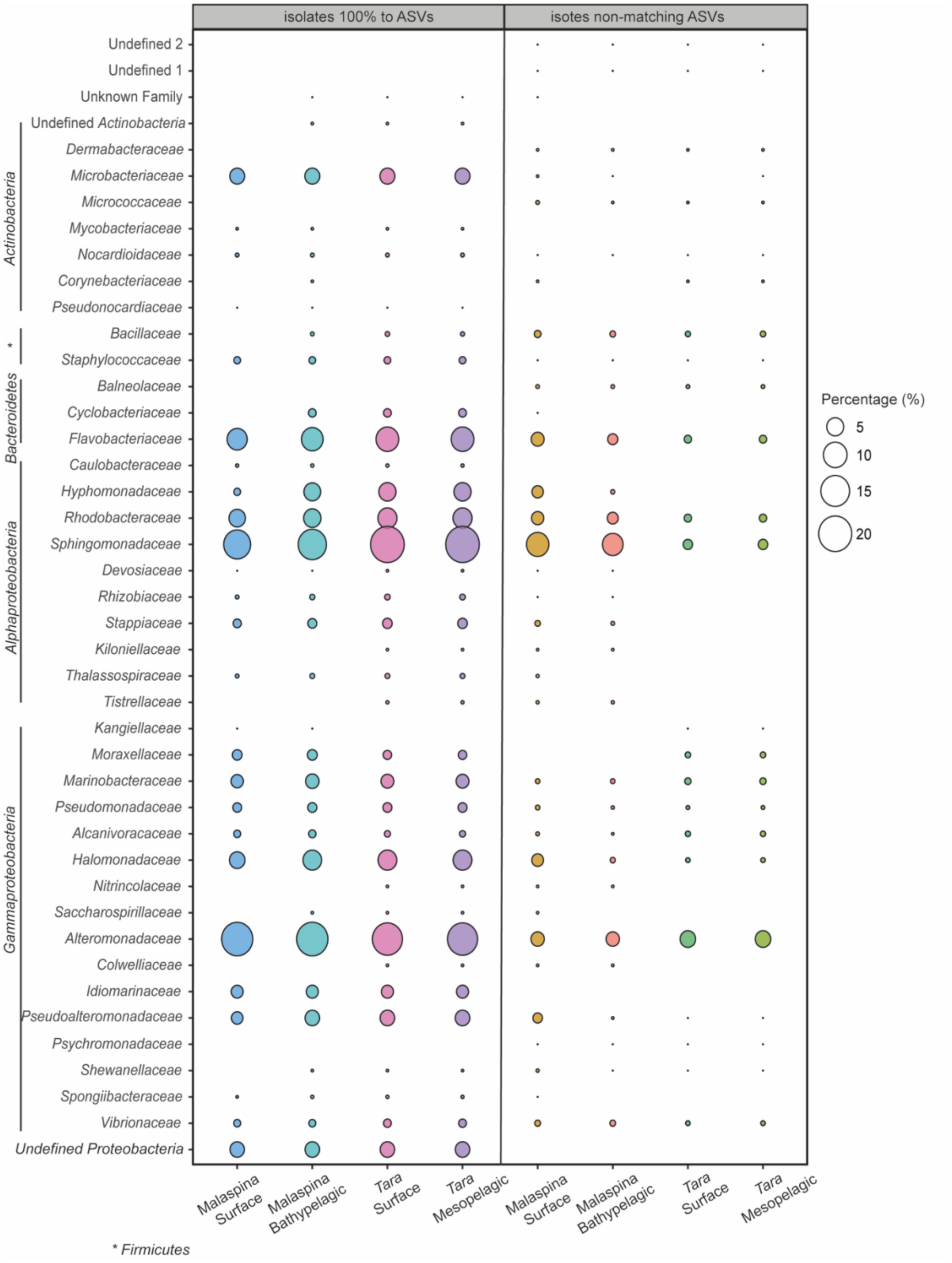

**Figure S3. Taxonomic differences at the family level between those isolates that are 100% identical to ASVs in the Malaspina Surface, Malaspina Bathypelagic, *Tara* Surface and *Tara* Mesopelagic datasets.** The first four columns represent those isolates 100% identical to ASVs, and the last four columns represent those isolates that did not match with any ASV. The size of the dots indicates the percentage of isolates in each family from the total isolates of the MARINHET_v2 collection.

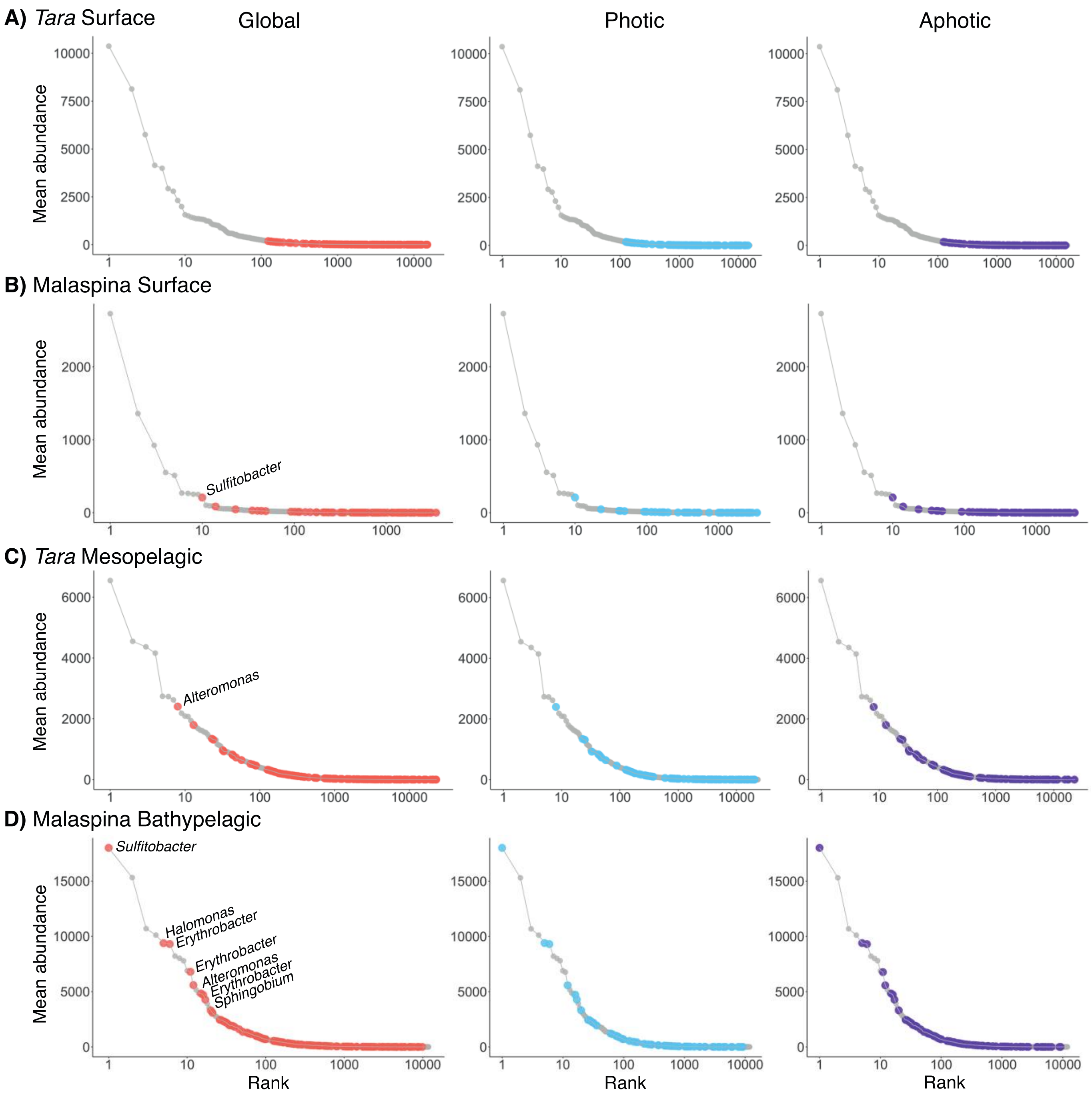

**Figure S4**. **Rank plots showing the ASVs recruited by photic and aphotic ocean isolates by each different dataset. (A)** *Tara* Surface. **(B)** Malaspina Surface. **(C)** *Tara* Mesopelagic. **(D)** Malaspina Bathypelagic. Coloured dots indicates which ASVs are 100% identical to at least one isolate in the comparisons made with all photic and aphotic depth isolates (global), and separately with the photic and aphotic datasets: grey, nonisolated zOTUs; orange, blue or purple, ASVs identical to isolates. Taxonomic affiliation is indicated for the abundant (>1 % abundance) ASVs that are identical to isolates in the global rank plot.

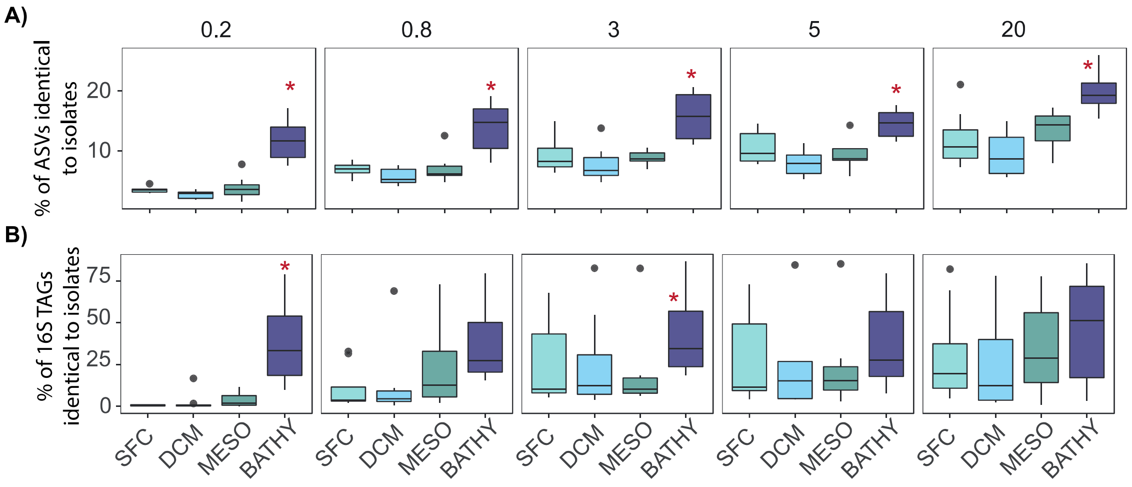

**Figure S5**. **Boxplots comparing different size fractions in the Malaspina profiles showing: (A)** percent of ASVs that were 100% identical to at least one isolate per size fractions and depth, and **(B)** percentages of 16S TAGs (reads) that were 100% identical to at least one isolate per size fractions and depth. SFC: surface, DCM: deep chlorophyll maximum, MESO: mesopelagic, and BATHY: bathypelagic. 0.2: free-living bacteria, 0.8: bacteria attached to small particles, and 3.0-20: bacteria attached to larger particles.. Significative differences between layers (Kruskal-Wallis, P.value from $1.1 \times 10^{-8}$ to $4.7 \times 10^{-12}$) are indicated by red asterisks.

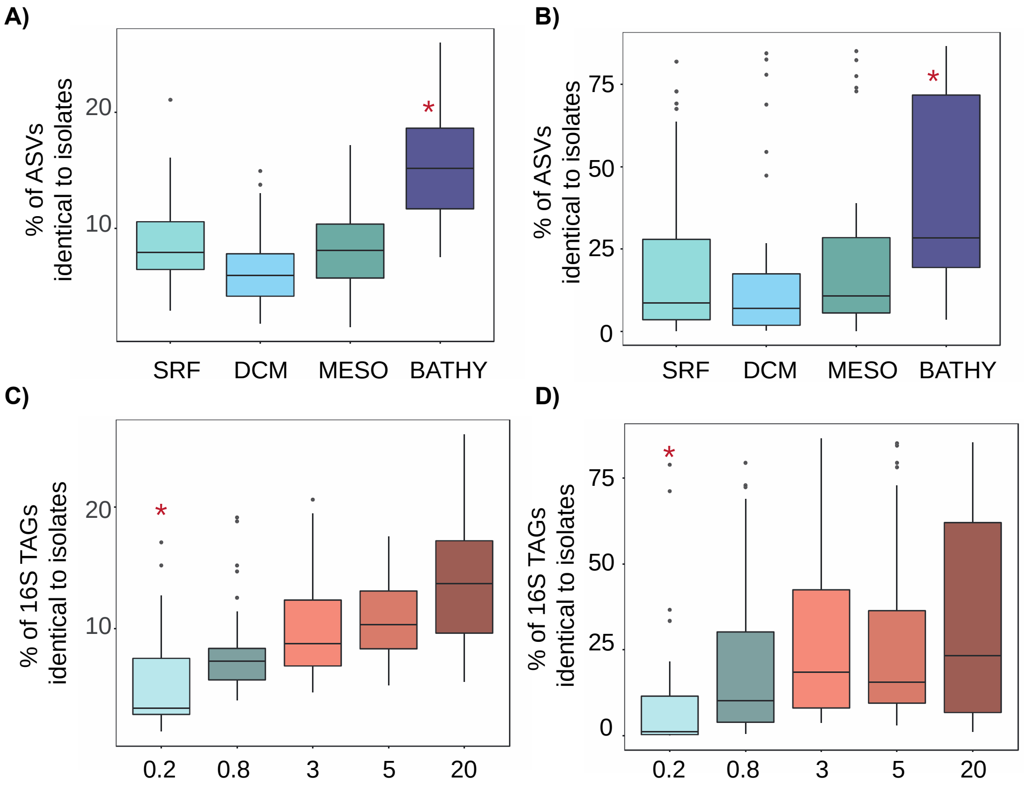

**Figure S6. Boxplots comparing different size fractions in the Malaspina profiles showing: (A,B)** percent of ASVs and 16S TAGs, respectively, that were 100% identical to at least one isolate per depth, and **(C,D)** percent of ASVs and 16S TAGs, respectively, that were 100% identical to at least one isolate per size fraction. SFC: surface, DCM: deep chlorophyll maximum, MESO: mesopelagic, and BATHY: bathypelagic. 0.2: free-living bacteria, 0.8: bacteria attached to small particles, and 3-20: bacteria attached to larger particles. Sizes of the fractions are in μm. Significative differences between layers or size fractions (Kruskal-Wallis, P.value <0.05) are indicated by red asterisks.

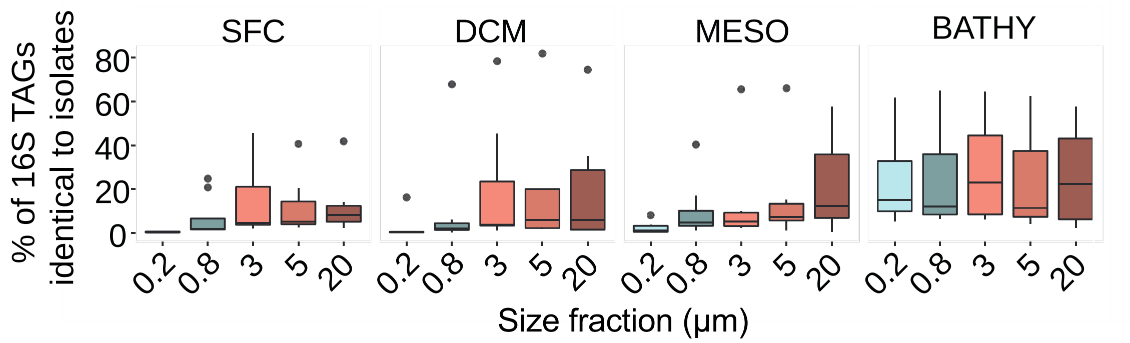

**Figure S7. Distribution of bathypelagic abundant ASVs (>1 % of the reads) in all layers and size fractions**. The values represent the mean abundance of reads across layers and fractions. SFC: surface, DCM: deep chlorophyll maxima, MESO: mesopelagic, and BATHY: bathypelagic. 0.2: free-living bacteria, 0.8: bacteria attached to small particles, and 3-20: bacteria attached to large particles.

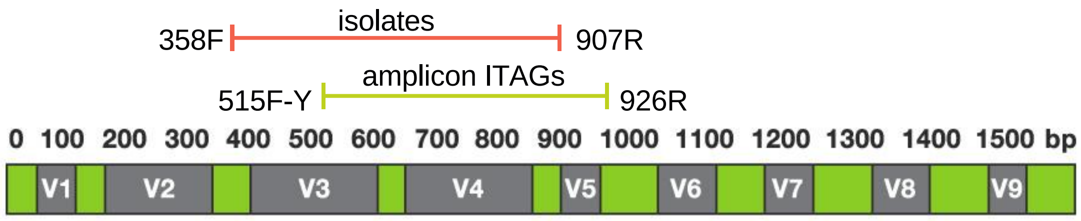

**Figure S8. Graphic representation of the 16S rRNA gene and the regions covered by isolates (orange) and amplicon 16S iTAGs (yellow).** Forward and reverse primers used for 16S rRNA amplifications in isolates and amplicon TAGs are indicated at the left and right sides of the lines.

#### Headings Supplementary Tables

**Table S1.** Summary of the total reads (16S TAGs), total OTUs (97%) and ASVs (100%), lowest number of reads and reads after rarification for each *Tara* Oceans and Malaspina Expedition amplicon 16S TAGs datasets. OTUs at 97% sequence similarity were obtained using the UPARSE algorithm of USEARCH v.10.0.240 (Edgar, 2010) just for comparative purposes.

| Dataset | UPARSE 97% | | | | | UNOISE 100% | | | | |
|---|---|---|---|---|---|---|---|---|---|---|
| | Total reads | Total OTUs | Lowest number of reads | Reads after rarification | OTUs after Rarification | Total reads | Total ASVs | Lowest number of reads | Reads after rarification | zOTUs after Rarification |
| *Malaspina Surface* | 6.16E+06 | 11685 | 12049 | 1373586 | 4975 | 6.34E+06 | 3810 | 12115 | 1381110 | 3528 |
| *Malaspina Bathypelagic* | 3.96E+07 | 26311 | 431835 | 17705235 | 19591 | 3.98E+07 | 11963 | 434069 | 17796829 | 11820 |
| *Tara Surface* | 3.46E+07 | 82671 | 140076 | 11206080 | 37064 | 3.64E+07 | 17270 | 160849 | 12867920 | 15426 |
| *Tara Mesopelagic* | 1.84E+07 | 79691 | 230364 | 8984196 | 48476 | 1.90E+07 | 24435 | 236694 | 9231066 | 23274 |

**Table S2.** Proportions of ASVs 100 % similarity and OTUs at 97 % identical to isolates for the three levels of comparisons: global, photic and aphotic zones, and per stations used for isolation.

| Dataset | Comparison | | OTUs identical to isolates (%) | | 16S TAGs identical to isolates (%) | |
|---|---|---|---|---|---|---|
| | | | *OTUs 97%* | *ASVs 100%* | *OTUs 97%* | *ASVs 100%* |
| **Tara Surface** | All isolates | | 3.8 | 2.3 | 4.8 | 1.6 |
| | Photic isolates | | 2.6 | 1.4 | 3.4 | 1.3 |
| | Aphotic isolates | | 2.4 | 1.5 | 2.7 | 1.3 |
| | Individual Surface Samples | ST39 | 0.3 | 0.2 | 1.7 | 1.4 |
| | | ST67 | 0.3 | 0.3 | 0.2 | 0.07 |
| | | ST72 | 0.6 | 0.4 | 1.8 | 1.2 |
| | | ST76 | 0.3 | 0.4 | 2.1 | 1.9 |
| | | ST151 | 0.8 | 0.4 | 1.7 | 0.9 |
| | | ST163 | 0.6 | 0.2 | 0.2 | 0.08 |
| | | ST84 | 0.5 | 0.3 | 0.08 | 0.07 |
| | | ST85 | 2.1 | 1.2 | 0.09 | 0.03 |
| **Tara Mesopelagic** | All isolates | | 2.6 | 1.7 | 9.8 | 8.5 |
| | Photic isolates | | 1.5 | 0.9 | 7.5 | 5.6 |
| | Aphotic isolates | | 1.6 | 1.1 | 8.8 | 7.8 |
| | Individual Mesopelagic Samples | ST102 | 0.2 | 0.2 | 11.2 | 9.2 |
| | | ST111 | 0.6 | 0.3 | 8.2 | 6 |
| | | ST138 | 0.7 | 0.4 | 9.5 | 6.1 |
| **Malaspina Surface** | All isolates | | 8.2 | 4.5 | 6.2 | 4.8 |
| | Photic isolates | | 7 | 2.7 | 5.6 | 3.3 |
| | Aphotic isolates | | 6.6 | 4.1 | 5.2 | 4.5 |
| **Malaspina Bathypelagic** | All isolates | | 3.1 | 2.4 | 35.7 | 27.9 |
| | Photic isolates | | 2 | 1.2 | 29.1 | 18.4 |
| | Aphotic isolates | | 2.1 | 1.8 | 32.4 | 26 |
| | Individual Bathypelagic Samples | ST10 | 0.2 | 0.1 | 14.1 | 10.4 |
| | | ST17 | 0.6 | 0.3 | 26.3 | 21.6 |
| | | ST23 | 1.2 | 0.5 | 32.2 | 28.9 |
| | | ST32 | 0.9 | 0.3 | 29.3 | 18.2 |
| | | ST43 | 0.3 | 0.2 | 19.9 | 16.4 |

**Table S3.** Proportions of isolates 100 % identical to ASVs per dataset, and proportions of those that did not match with any ASV.

| | | | | |
|---|---|---|---|---|
| Isolates 100% identical to ASVs (%) | 1445 (72.1%) | 1639 (81.8%) | 1787 (89.2%) | 1784 (89%) |
| Isolates with 1 ASV hit (%) | 1420 (70.9%) | 1613 (80.5%) | 1763 (87.9%) | 1762 (87.95) |
| Isolates with 2 ASVs hit (%) | 26 (1.3%) | 27 (1.3%) | 24 (1.2%) | 24 (1.2%) |
| Non-matching isolates (%) | 559 (27.9%) | 365 (18.2%) | 217 (10.8%) | 220 (10.9%) |

**Table S4.** Taxonomic classification of the different ASVs that were 100% identical to isolates found in *Tara* Oceans and Malaspina Expedition amplicon 16S iTAGs datasets.

*** file in Excel document**

**Table S5.** Proportion of 16S TAGs (reads) identical to isolates at 100 % similarity in the Malaspina size fraction dataset per layer and size fraction. SFC: surface, DCM: deep chlorophyll maximum, MESO: mesopelagic; BATHY: bathypelagic; sd, standard deviation; IQR, interquartile range.

| **Proportion of 16s TAGs identical to isolates per layer and size fraction** | | | | | | |
|---|---|---|---|---|---|---|
| DEPTH | FILTER | stations_cou | mean | sd | median | IQR |
| SFC | 0.2 | 8 | 0.0 | 0.0 | 0.0 | 0.0 |
| SFC | 0.8 | 8 | 0.1 | 0.1 | 0.0 | 0.1 |
| SFC | 3 | 8 | 0.3 | 0.3 | 0.2 | 0.4 |
| SFC | 5 | 8 | 0.3 | 0.3 | 0.2 | 0.3 |
| SFC | 20 | 8 | 0.3 | 0.3 | 0.2 | 0.3 |
| DCM | 0.2 | 8 | 0.0 | 0.1 | 0.0 | 0.0 |
| DCM | 0.8 | 8 | 0.1 | 0.2 | 0.1 | 0.0 |
| DCM | 3 | 8 | 0.3 | 0.3 | 0.1 | 0.2 |
| DCM | 5 | 8 | 0.2 | 0.3 | 0.2 | 0.2 |
| DCM | 20 | 8 | 0.2 | 0.3 | 0.1 | 0.2 |
| MESO | 0.2 | 8 | 0.0 | 0.0 | 0.0 | 0.1 |
| MESO | 0.8 | 8 | 0.2 | 0.2 | 0.1 | 0.3 |
| MESO | 3 | 8 | 0.2 | 0.3 | 0.2 | 0.2 |
| MESO | 5 | 8 | 0.2 | 0.3 | 0.2 | 0.1 |
| MESO | 20 | 8 | 0.3 | 0.3 | 0.2 | 0.4 |
| BATHY | 0.2 | 7 | 0.4 | 0.3 | 0.3 | 0.4 |
| BATHY | 0.8 | 7 | 0.4 | 0.3 | 0.3 | 0.3 |
| BATHY | 3 | 7 | 0.4 | 0.3 | 0.3 | 0.3 |
| BATHY | 5 | 7 | 0.4 | 0.3 | 0.3 | 0.4 |
| BATHY | 20 | 7 | 0.5 | 0.3 | 0.5 | 0.5 |

**Table S6.** Proportion of ASVs identical to isolates at 100 % similarity in the Malaspina size fraction dataset per layer and size fraction. SFC: surface, DCM: deep chlorophyll maximum, MESO: mesopelagic; BATHY: bathypelagic; sd, standard deviation; IQR, interquartile range.

| **Proportion of ASVs identical to isolates per layer and size fraction** | | | | | | |
|---|---|---|---|---|---|---|
| DEPTH | FILTER | count | mean | sd | median | IQR |
| SFC | 0.2 | 8 | 0.0 | 0.0 | 0.0 | 0.0 |
| SFC | 0.8 | 8 | 0.1 | 0.0 | 0.1 | 0.0 |
| SFC | 3 | 8 | 0.1 | 0.0 | 0.1 | 0.0 |
| SFC | 5 | 8 | 0.1 | 0.0 | 0.1 | 0.0 |
| SFC | 20 | 8 | 0.1 | 0.0 | 0.1 | 0.0 |
| DCM | 0.2 | 8 | 0.0 | 0.0 | 0.0 | 0.0 |
| DCM | 0.8 | 8 | 0.1 | 0.0 | 0.1 | 0.0 |
| DCM | 3 | 8 | 0.1 | 0.0 | 0.1 | 0.0 |
| DCM | 5 | 8 | 0.1 | 0.0 | 0.1 | 0.0 |
| DCM | 20 | 8 | 0.1 | 0.0 | 0.1 | 0.0 |
| MESO | 0.2 | 8 | 0.0 | 0.0 | 0.0 | 0.0 |
| MESO | 0.8 | 8 | 0.1 | 0.0 | 0.1 | 0.0 |
| MESO | 3 | 8 | 0.1 | 0.0 | 0.1 | 0.0 |
| MESO | 5 | 8 | 0.1 | 0.0 | 0.1 | 0.0 |
| MESO | 20 | 8 | 0.1 | 0.0 | 0.1 | 0.0 |
| BATHY | 0.2 | 7 | 0.1 | 0.0 | 0.1 | 0.1 |
| BATHY | 0.8 | 7 | 0.2 | 0.0 | 0.2 | 0.1 |
| BATHY | 3 | 7 | 0.2 | 0.0 | 0.2 | 0.1 |
| BATHY | 5 | 7 | 0.2 | 0.0 | 0.1 | 0.0 |
| BATHY | 20 | 7 | 0.2 | 0.0 | 0.2 | 0.0 |

**Table S7.** Summary of the proportions of ASVs and 16S TAGs identical (100% similarity) to isolates after removing *Cyanobacteria* (A) or *Archaea* (B) from the ASV-abundance tables in photic and aphotic zone *Tara* Oceans and Malaspina Expedition datasets.

| A. ASVs Iidentucal to isolates without cyanos | | | |
|---|---|---|---|
| **Dataset** | **Comparison** | **% ASVs** | **% 16S TAGs** |
| Tara Surface | All isolates | 2.9 | 2.9 |
| Tara Meso | All isolates | 2 | 10.9 |
| Malaspina Surface | All isolates | 6.1 | 10.1 |
| Malaspina Bathypelagic | All isolates | 2.7 | 29.1 |

| B. ASVs identical to isolates without Archaea | | | |
|---|---|---|---|
| **Dataset** | **Comparison** | **% ASVs** | **% 16S TAGs** |
| Tara Surface | All isolates | 2.23 | 1.61 |
| Tara Meso | All isolates | 1.9 | 11.63 |
| Malaspian Surface | All isolates | 4.64 | 4.96 |
| Malaspina Bathypelagic | All isolates | 2.60 | 31.43 |

**Table S8.** Summary of the rRNA average copy number for each of the genera identified in the MARINHET culture collection. Values extracted from: https://rrndb.umms.med.umich.edu/.

| Phyllum | Genera | Número de rrna operon | |
|---|---|---|---|
| | | mean | sd |
| *Actinobacteria* | *Corynebacterium* | 4.3 | 0.8 |
| | *Mycobacterium* | 1.1 | 0 |
| | *Microbacterium* | 2 | 0.4 |
| | *Nocardioides* | 2.4 | 0.8 |
| | *Pseudonocardia* | 3.7 | 0.8 |
| | *Undefined* | NA | |
| *Bacteroidetes* | *Algoriphagus* | 3 | 0 |
| | *Roseivirga* | 1 | 0 |
| | *Croceibacter* | 2 | 0 |
| | *Dokdonia* | 3 | 0 |
| | *Gramella* | 3 | 0 |
| | *Joostella* | NA | |
| | *Leeuwenhoekiella* | 3 | 0 |
| | *Marixanthomonas* | 2 | 0 |
| | *Mesoflavibacter* | 2 | 0 |
| | *Mesonia* | NA | |
| | *Muricauda* | 2 | 0 |
| | *Polaribacter* | 3.9 | |
| | *Salegentibacter* | 3 | 0 |
| | *Vitellibacter* | 2 | 0 |
| | *Winogradskyella* | 2.6 | 1.3 |
| | *Zunongwangia* | 3 | 0 |
| | *Flavobacteriaceae* | 4.6 | 2.9 |
| *Firmicutes* | *Bacillus* | 8.7 | 3.3 |
| | *Staphylococcus* | 5.7 | 0.7 |
| *Alphaproteobacteria* | *Acuticoccus* | 2 | 0 |
| | *Brevundimonas* | 2 | 0.4 |
| | *Hyphomonas* | 1 | 0 |
| | *Maricaulis* | 2 | 0 |
| | *Oceanicaulis* | 1.3 | 0.4 |
| | *Ponticaulis* | 1.2 | 0.4 |
| | *Devosia* | 2.4 | 1.4 |
| | *Aurantimonas* | 2 | 0 |
| | *Martelella* | 2.5 | |
| | *Nitratireductor* | 1.7 | 0.5 |
| | *Pseudohoeflea* | 1.9 | 0.5 |
| | *Labrenzia* | 2.9 | 0.3 |
| | *Stappia* | 2.8 | 0.4 |
| | *Loktanella* | 2.7 | 1.2 |
| | *Paracoccus* | 2.7 | 0.5 |
| | *Pseudooceanicola* | 2 | 0 |
| | *Pseudoruegeria* | 2.7 | 1.2 |
| | *Roseovarius* | 1.2 | 0.4 |
| | *Shimia* | 2.7 | 1.2 |
| | *Sulfitobacter* | 2.5 | 1.2 |
| | *Thiobacimonas* | NA | |
| | *Thioclava* | 3 | 0 |
| | *Thalassospira* | 4 | 1.1 |
| | *Tistilla* | NA | |
| | *Tistrella* | 4 | 0 |
| | *Altererythrobacter* | 1.3 | 0.4 |
| | *Erythrobacter* | 1.7 | 1 |
| | *Novosphingobium* | 2.8 | 1.3 |
| | *Sphingobium* | 2.9 | 1 |
| | *Sphingopyxis* | 1.1 | 0.2 |
| | *Sphingimonadaceae* | 2.1 | 1 |
| | *Rhodobacteraceae* | 2.8 | 1.1 |
| *Gammaproteobacteria* | *Aestuariibacter* | 6.2 | 2.5 |
| | *Agaribacter* | 6.2 | 2.5 |
| | *Alteromonas* | 7.1 | 2.2 |
| | *Paraglaciecola* | 5.3 | 0.4 |
| | *Colwellia* | 6.7 | 1.2 |
| | *Idiomarina* | 4 | 0 |
| | *Marinobacter* | 3.3 | 0.9 |
| | *Pseudoalteromonas* | 8.7 | 1.1 |
| | *Shewanella* | 8.8 | 1.6 |
| | *Spongiibacter* | 2.3 | 0.4 |
| | *Alcanivorax* | 2.4 | 0.5 |
| | *Chromohalobacter* | 5 | 0 |
| | *Cobetia* | 6.9 | 0.3 |
| | *Halomonas* | 5.3 | 1.1 |
| | *Kushneria* | 4 | 0 |
| | *Salinicola* | 5 | 0 |
| | *Marinobacterium* | 4.8 | 1.1 |
| | *Kangiella* | 2 | 0 |
| | *Oleispira* | 5.6 | 2 |
| | *Acinetobacter* | 6.2 | 0.5 |
| | *Psychrobacter* | 4.4 | 0.9 |
| | *Pseudomonas* | 4.9 | 1.2 |
| | *Alteromonadaceae* | 6.2 | 2.5 |
| | *Vibrionaceae* | 10 | 3.3 |
| | *Undefined* | NA | |
| *AVERAGE rRNA copy number* | | 3.515789474 | -- |

**Table S9.** Mean abundance of the most abundant ASVs (>1% reads) in the bathypelagic (BATHY) samples extracted from the Malaspina size fraction dataset and their respective mean abundances in the surface (SFC), deep chlorophyll maxima (DCM) and mesopelagic (MESO) samples per size fraction.

| Proportion of bathypelagic abundant ASVs identical to isolates per layer and size fraction | | | | |
|---|---|---|---|---|
| FILTER | DEPTH | count | mean abundance | % mean abundance |
| 0.2 | BATHY | 7 | 0.24 | 7.62 |
| 0.8 | BATHY | 7 | 0.25 | 7.84 |
| 3 | BATHY | 7 | 0.28 | 9.10 |
| 5 | BATHY | 7 | 0.24 | 7.63 |
| 20 | BATHY | 7 | 0.26 | 8.24 |
| 0.2 | DCM | 8 | 0.02 | 0.73 |
| 0.8 | DCM | 7 | 0.12 | 3.71 |
| 3 | DCM | 8 | 0.19 | 6.17 |
| 5 | DCM | 5 | 0.22 | 7.14 |
| 20 | DCM | 6 | 0.21 | 6.59 |
| 0.2 | MESO | 8 | 0.02 | 0.69 |
| 0.8 | MESO | 8 | 0.10 | 3.25 |
| 3 | MESO | 6 | 0.15 | 4.85 |
| 5 | MESO | 8 | 0.15 | 4.75 |
| 20 | MESO | 7 | 0.22 | 7.10 |
| 0.2 | SFC | 6 | 0.00 | 0.10 |
| 0.8 | SFC | 8 | 0.07 | 2.19 |
| 3 | SFC | 7 | 0.14 | 4.63 |
| 5 | SFC | 7 | 0.12 | 3.86 |
| 20 | SFC | 8 | 0.12 | 3.78 |

**Table S10.** Characteristics of the different samples from where isolates of marine heterotrophic bacteria were obtained. 'Non-redundant isolates' are the isolates remaining after duplicates (100% identical in their partial 16S rRNA gene) are removed.

| Expedition | Station | Ocean | Latitude | Longitude | Depth (m) | *In situ* temperature (ºC) | Nº of sequenced isolates | Nº of non-redundant isolates | Cfu/ml | Cells/ml |
|---|---|---|---|---|---|---|---|---|---|---|
| *Tara* Oceans | ST 39 | Indian Ocean | 19º 2.24' N | 64º 29.24' E | 5.5 | 26.2 | 109 | 25 | 2.83E+03 | 9.70E+05 |
| | ST 39 | Indian Ocean | 18º 35.2' N | 66º 28.22' E | 25 | 26.8 | 243 | 53 | NA | 9.50E+05 |
| | ST 39 | Indian Ocean | 18º 43.12' N | 66º 21.3' E | 268.2 | 15.6 | 88 | 18 | NA | 9.70E+05 |
| | ST 67 | South Atlantic | 32º 17.31' S | 17º 12.22' E | 5 | 12.8 | 115 | 49 | 2.24E+03 | 1.90E+06 |
| | ST 72 | South Atlantic | 8º 46.44' S | 17º 54.36' W | 5 | 25 | 71 | 33 | 1.01E+04 | 6.90E+05 |
| | ST 76 | South Atlantic | 20º 56.7' S | 35º 10.49' W | 5 | 23.3 | 89 | 27 | 1.06E+03 | 6.10E+05 |
| | ST 84 | Southern Ocean | 60º 13.4' S | 60º 38.51' W | 5.9 | 1.8 | 10 | 8 | 1.28E+02 | 2.65E+05 |
| | ST 85 | Southern Ocean | 62º 2.19' S | 49º 31.44' W | 5.9 | 0.7 | 126 | 30 | 4.28E+02 | 4.11E+05 |
| | ST 85 | Southern Ocean | 62º 2.19' S | 49º 31.44' W | 87.4 | -0.8 | 13 | 10 | 7.00E+01 | 2.02E+05 |
| | ST 102 | Pacific Ocean | 5º 16.12' S | 85º 13.12' O | 475.6 | 9.2 | 97 | 15 | 2.35E+03 | 1.60E+05 |
| | ST 111 | Pacific Ocean | 16º 57.36' S | 100º 39.36'O | 347.1 | 10.9 | 98 | 35 | 1.66E+03 | 6.50E+04 |
| | ST 138 | Pacific Ocean | 6º 22.12' N | 103º 4.12' O | 444.9 | 8.2 | 91 | 34 | 1.12E+03 | 1.30E+05 |
| | ST 151 | North Atlantic | 36º 10.17' N | 29º 1.23' W | 5 | 17.3 | 76 | 33 | 2.28E+03 | 4.40E+05 |
| | ST 163 | Arctic Ocean | 76º 10.57' N | 1º 23.31' E | 5 | 1.9 | 18 | 7 | 8.17E+01 | 1.80E+05 |
| | ST 175 | Arctic Ocean | 79º 13.24' N | 66º 20.37' E | 5 | 1.4 | 3 | 3 | 2.50E+01 | 4.21E+05 |
| | ST 201 | Arctic Ocean | 74º 17.23º N | 85º 48' W | 5 | -1.3 | 1 | 1 | NA | 5.88E+02 |
| ATP 09 | AR_1 | Arctic Ocean | 78º 20.00' N | 15º 00.00' E | 2 | 6.2 | 13 | 9 | NA | NA |
| | AR_2 | Arctic Ocean | 76º 28.65' N | 28º 00.62' E | 25 | -1.2 | 20 | 9 | NA | NA |
| Malaspina | ST 10 | North Atlantic | 21º 33.36' N | 23º26' W | 4002 | 2.04 | 20 | 9 | 5.00E+01 | 3.30E+04 |
| | ST 17 | South Atlantic | 3º 1.48' S | 27º 19.48' W | 4002 | 1.74 | 93 | 24 | 7.73E+02 | 3.00E+04 |
| | ST 23 | South Atlantic | 15º 49.48' S | 33º 24.36' W | 4003 | 1.45 | 94 | 39 | 4.38E+02 | 1.20E+04 |
| | ST 32 | South Atlantic | 26º 56.8' S | 21º 24' W | 3200 | 2.5 | 62 | 20 | 1.90E+02 | 1.80E+04 |
| | ST 33 | South Atlantic | 27º 33.2' S | 18º 5.4' W | 3904 | 1.7 | 28 | 13 | 3.70E+01 | 1.50E+04 |
| | ST 43 | South Atlantic | 32º 48.8' S | 12º 46.2' E | 4000 | 1.2 | 46 | 19 | 3.00E+01 | 3.80E+04 |
| MIFASOL | ST 8 | NW Mediterranean | 40º 38.41' N | 2º 50' E | 2000 | 13.2 | 245 | 62 | 5.54E+02 | 2.20E+04 |
| BBMO | BBMO | NW Mediterranean | 41º 40' N | 2º 48' E | 5 | 17.71 | 134 | 62 | NA | 6.70E+05 |

**Table S11.** Culture media and incubation conditions used for each sample. Positive signs indicate which media were used. RT: room temperature.

| Cruise | Depth and station | Tº | Zobell agar | Marine agar 2216 | modified marine agar 2216 |
|---|---|---|---|---|---|
| *Tara* Oceans | 5 m (stations 67, 72, 76, 151) | RT | - | + | - |
| | 5 m (stations 84, 85, 163, 175, 201) | RT/4ºC | - | - | + |
| | 25 m (station 85) | RT/4ºC | - | - | + |
| | 25 m (station 39) | RT | + | - | - |
| | OMZ (stations 39, 102, 111, 138) | RT | + | - | - |
| ATP 09 | 5 m | 4ºC | + | - | - |
| | 25 m | 4ºC | + | - | - |
| Malaspina | 4000 m (stations 10, 17, 23) | RT | - | + | - |
| | 4000 m (stations 32, 33, 43) | RT/4ºC | + | + | + |
| Mifasol | 2000 m | RT/12ºC | + | + | + |
| BBMO | 5 m | RT | + | + | + |
